# Haplotype-Based Models Improve Sweep Detection in Ancient Populations with Complex Demography

**DOI:** 10.64898/2026.05.08.723766

**Authors:** Abigail N. Sequeira, Zachary A. Szpiech, Christian D. Huber

## Abstract

Identifying signatures of positive selection in humans is complicated by demographic processes such as bottlenecks, migration and admixture, all of which can distort or obscure the genomic patterns produced by selective sweeps. Ancient DNA offers a direct window into past allele and haplotype frequencies, yet most sweep scans in ancient populations rely on allele-frequency or site frequency spectrum (SFS) summaries, with limited use of haplotype-based approaches. Here, we evaluate the performance of haplotype and SFS-based methods for detecting selective sweeps under demographic scenarios that reflect the complex history of ancient and modern Europeans. We extend the haplotype-based likelihood framework saltiLASSI to accommodate pseudohaploid ancient genomes, enabling the use of truncated haplotype frequency spectra and their spatial decay to detect sweeps without requiring phased data. Using forward-in-time simulations, we examine sweeps of varying ages, two pulses of admixture with different source proportions, and cases where selection continues or ceases after admixture. We compare saltiLASSI to a widely used SFS-based approach (SweepFinder2). Our results show that haplotype-based likelihood models retain higher power than SFS methods in admixed populations, particularly when sweep haplotypes are introduced through migration or when selection has not had sufficient time to regenerate a clear SFS signature after admixture. These findings highlight the promise of haplotype-based inference for ancient DNA and demonstrate how model-based approaches can improve the detection of historical selective sweeps in populations with complex demographic histories.

## Introduction

Natural selection leaves characteristic patterns in the genome that can be used to infer past evolutionary processes. A classic example is the selective sweep, in which a beneficial mutation rises rapidly in high frequency along with any “hitch-hiking” linked variants (Smith and Haigh, 1974). Such sweeps generate a reduction in local nucleotide diversity (Kim and Stephan, 2002), elevated linkage disequilibrium (LD) (Kim and Nielsen, 2004), and distortions of the site frequency spectrum (SFS) (Fay and Wu, 2000; Kim, 2006). The extent of these signals depends on recombination rate, with higher recombination rate producing narrower sweep footprints and lower recombination producing broader ones (Begun and Aquadro, 1992). Haplotype structure provides additional information, as regions surrounding a selected allele show increased haplotype homozygosity and stronger associations among nearby variants (Sabeti *et al*., 2002; Kim and Nielsen, 2004). These signatures erode over time as new mutations, recombination, and drift restore diversity levels and break down LD around the selected site.

Although these genomic features underpin many approaches to sweep detection, their interpretation can be complicated by demographic history (Harris, Sackman, *et al*., 2018; Schneider *et al*., 2021). Recent and severe bottlenecks increase false positives due to an excess of rare variants that mimic the genomic pattern of a selective sweep (Przeworski, 2002; Jensen *et al*., 2005). Moreover, human populations have become less differentiated due to the high levels of migration and admixture during the Holocene (Lazaridis *et al*., 2016; Williams *et al*., 2024).

Admixture is known to obscure sweep signals in both ancient and modern populations by reducing the sweep haplotype frequency (Souilmi *et al*., 2022; Pandey *et al*., 2024), particularly if not enough time has passed for the sweep to increase in frequency again, or selection is no longer acting on the trait (Souilmi *et al*., 2022). Alternatively, admixture could introduce an adaptive haplotype into the receiving population and become the dominant haplotype; such is evidenced with the introduction of the lactase persistence allele at *LCT* from the Steppe pastoralists into Europeans around 3,000 years ago (Allentoft *et al*., 2015).

Ancient DNA provides direct snapshots of populations before and after major demographic events, helping to recover sweep signals otherwise obscured in modern populations (Mallick *et al*., 2024). Ancient DNA studies have relied on allele frequency-based tests (Mathieson *et al*., 2015; Ye *et al*., 2017; Akbari *et al*., 2026) or, more rarely, on SFS approaches such as SweepFinder2 (SF2) (Souilmi *et al*., 2022). In contrast, haplotype-based methods have seen limited application in ancient populations (Martiniano *et al*., 2017; Childebayeva *et al*., 2022; Pandey *et al*., 2024), in part because many require phased genotypes (Voight *et al*., 2006; Ferrer-Admetlla *et al*., 2014; Garud *et al*., 2021; Szpiech *et al*., 2021). Recent developments now permit haplotype-based inference from unphased data (Harris, Garud, *et al*., 2018; DeGiorgio and Szpiech, 2022; Szpiech, 2024), opening the door to a broader use in ancient DNA. These methods capture information about linkage disequilibrium and haplotype structure that is not available from SFS-based tests (Kim and Nielsen, 2004), and therefore may retain greater power to detect recent selective sweeps and sweeps in populations with histories of admixture (Ferrer-Admetlla *et al*., 2014; Harris and DeGiorgio, 2020).

To date, only three studies have applied haplotype-based methods to ancient DNA (Martiniano *et al*., 2017; Childebayeva *et al*., 2022; Pandey *et al*., 2024). Martiniano et al. (Martiniano *et al*., 2017) examined the extended haplotype homozygosity (EHH) around known sweep-associated SNPs in Europeans, but did not perform a genome-wide scan. Childebayeva et al. (2022) tested the cross population extended haplotype homozygosity (XP-EHH) test on imputed ancient data. Pandey et al. (Pandey *et al*., 2024) used the haplotype statistic G12 on simulated ancient data under low and continuous migration scenarios, showing that haplotype-based tests can be applied to pseudohaploidized genomes. However, their window-based approach does not incorporate the full haplotype frequency spectrum (HFS) or broader haplotype structure along the genome. Here, we extend the haplotype-based likelihood framework saltiLASSI (SL) (DeGiorgio and Szpiech, 2022) for use with ancient DNA. Unlike other available methods, SL models the spatial genomic decay of the truncated HFS to identify sweep footprints. We evaluate its performance under a range of sweep ages, two admixture pulses with different source contributions and timings, and scenarios in which selection either persists or ceases after admixture. This simulation framework more closely reflects the demographic history of modern Europeans than in previous work and supports saltiLASSI as a promising approach for detecting selective sweeps in ancient genomes and for characterizing changes in haplotype structure through time.

## Methods

### Simulations

We simulated 200 hard selective sweeps for each population under the complex European demographic history described in Souilmi et al. (2022) using msprime v1.2.0 (Baumdicker et al. 2022) and SLiM v4.0.1 (Haller and Messer, 2023). Each simulation modeled a 5 Mb genomic region with a neutral mutation rate of 1.5×10⁻⁸ per base per generation and a recombination rate of 1.28×10⁻⁸ (Souilmi *et al*., 2022). A single beneficial mutation was introduced at the center of the region at a specified time point. Simulations in which the mutation was lost were restarted, and only realizations in which the mutation persisted were included in downstream analyses. For each demographic model, we simulated fully neutral scenarios (*s* = 0) as well as selective sweeps with selection coefficients *s* = 0.01, 0.02, and 0.1. We defined selection such that heterozygotes have fitness 1+*s* and homozygotes for the beneficial allele have fitness 1+2*s*. To reduce computational cost while preserving key population genetic properties, we applied a down-scaling scheme in which population sizes and times were divided by a factor of four, and selection coefficients and mutation and recombination rates were multiplied by the same factor.

### Admixture model

To simulate European demographic history, we followed the parameters and model used in Souilmi et al. (2022). We first generated 1,000 constant size burn-in simulations using msprime (Baumdicker *et al*., 2022), and for each forward simulation in SLiM4 (Haller and Messer, 2023) we randomly selected one burn-in as the ancestral starting point, just before the African and Eurasian split. The demographic model followed Souilmi et al. (2022) and included multiple splits giving rise to Anatolian early farmers, Western Hunter-Gatherers (WHG), Caucasus Hunter-Gatherers (CHG), and Ancient North Eurasians/Eastern Hunter-Gatherers (ANE/EHG). Steppe ancestry was modeled as admixture between CHG and ANE/EHG, and subsequent admixture events among Steppe, Anatolian, and WHG lineages produced Early European Farmers (EEF), Late Neolithic/Bronze Age populations (LNBA), and modern Europeans. We simulated three hard sweep start times, one before each major population split so that the beneficial mutation was either present in all three ancestral populations (Anatolia EF, WHG, and Steppe; 55 kya), in two of the ancestral populations (Anatolia EF and WHG; 44 kya) or in only one ancestral population (Anatolia EF; 36 kya). To assess the impact of admixture on the detectability of an ancestral sweep, we ran simulations in which selection was turned off at the time of admixture, allowing the sweep haplotype to be diluted by gene flow. We sampled individuals from the three ancestral populations, from ancient, admixed populations, and from simulated modern Europeans. Sample sizes for ancient groups were matched to those in our empirical ancient DNA dataset (see below).

To mimic ancient data, we pseudohaploidized the simulation VCFs by randomly picking one of the two alleles at each heterozygous site. To approximate SNP capture panels commonly used in ancient DNA studies, we selected one modern African and one modern European diploid genome from the simulation and ascertained SNPs as sites at which either individual was heterozygous. From these positions, we randomly selected 2,000 SNPs to approximate the number of variants targeted by common ancient capture arrays in a typical 5 Mb region. We then estimated the missing-data distribution from the empirical ancient dataset and, for each population separately, applied the corresponding distribution to the simulated SNP data to reproduce realistic levels of missingness.

### Detecting Selective Sweeps

We evaluated three methods for detecting selective sweeps: the SFS–based approach SweepFinder2 (SF2) (DeGiorgio *et al*., 2016) and the haplotype-based methods saltiLASSI (SL) (DeGiorgio and Szpiech, 2022) and G12 (Pandey *et al*., 2024). For saltiLASSI, we used a modified version of the original method (see below for details), whereas G12 was applied as described in Pandey et al. (2024). We ran SF2 by first estimating the background site frequency spectrum (SFS) from 600 neutral simulations for each population (*SweepFinder2-f neutral.freq.file SFS*). Then, we computed the composite likelihood ratio (CLR) every 1000 bp using the pre-calculated background SFS for each simulation (*SweepFinder2-lg 1000 sim.freq.file SFS Out.file*). We ran SL v1.2.2 using two different window sizes (51 and 101 SNPs) with the following parameters: *–winstep* = 1, *–filter-level* = 1, *–unphased*, *–k =* 5, *–salti*, *–max-hmiss = 0.99*, and *–max-lmiss = 0.99*. For each population, we calculated the null haplotype frequency spectrum (HFS) from the set of 600 neutral simulations and calculated likelihood ratios using the corresponding background HFS for each replicate. Together the whole saltiLASSI command followed:

*lassip –vcf pop.sim.vcf –k 5 –calc-spec –hapstats –unphased –max-hmiss 0.99 –max-lmiss 0.99 –match-tol 1 –winsize SNPsize –winstep 1 –filter-level 1 –pop pop_ID.txt –out pop.sim.out lassip –spectra pop.neutral.spectra –avg-spec –out pop.null.HFS lassip –spectra pop.sim.spectra –salti –null-spec pop.null.HFS –out pop.sim.out*

We applied G12 (Pandey *et al*., 2024) with the same window sizes (51 and 101 SNPs) and a step size of 1 to allow direct comparison with SL (*python H12_H2H1.py pop.sim.input.txt sampleSize-o pop.sim.out.txt-w SNPsize-j 1-d 1*). Both haplotype-based methods (SL and G12) were run on datasets before and after filtering monomorphic sites to evaluate how invariant positions influence their power to detect sweeps. For each simulation replicate, we recorded the maximum statistic value across the entire 5 Mb region (CLR for SF2, likelihood ratio for saltiLASSI, and G12). For each method and sweep scenario, the 1% false positive rate (FPR) cutoff was then defined as the 99th percentile of these maximum values across the 600 neutral simulations.

Power was calculated as the proportion of sweep simulations (*n* = 200) whose maximum statistic exceeded this threshold. We also conducted a pairwise chi-square test followed by a Benjamini and Hochberg correction for each sweep condition and population to determine whether any of the three methods perform significantly better than the other. The chi-square test results are reported in SI Tables 3, 5, 7, 10-11.

### Modifications to saltiLASSI for ancient DNA

saltiLASSI (SL) is a haplotype-based selection scan that builds on the *T* statistic introduced in LASSI (Harris and DeGiorgio, 2020). SL extends LASSI by modeling the spatial genomic pattern of the truncated HFS across adjacent windows, improving sensitivity to extended selection signals compared to single-window tests. While SL performs well on high-quality modern genomic data, its direct application to datasets with substantial missingness, such as ancient DNA, is challenging due to highly variable coverage across loci and individuals, where average coverage is often near or below 1× and pseudohaploidization is commonly applied. To address these limitations, we extended saltiLASSI by introducing tolerance-based clustering of pseudohaplotypes and user-defined filtering of loci (*--max-lmiss*) and haplotypes (*--max-hmiss*) based on missingness. Within each analysis window, pseudohaplotypes are compared pairwise, and those differing at fewer than or equal to a user-specified number of sites are grouped into a single haplotype class (*--match-tol*). Sequence similarity is quantified using Hamming distance, such that pseudohaplotypes that differ only by a small number of mismatches, arising from sequencing error or DNA decay, are treated as equivalent. The frequency of each clustered haplotype class is then computed from the number of contributing pseudohaplotypes and used to construct the truncated HFS underlying the SL statistic. This clustering strategy follows the general approach used for the H12 and H2/H1 statistics and reduces artificial inflation of rare haplotypes in low-coverage data (Garud *et al*., 2015). In addition to haplotype clustering, we extended saltiLASSI to flexibly filter loci (*--max-lmiss*) and pseudohaplotypes (*--max-hmiss*) based on their proportion of missing data. Loci or pseudohaplotypes exceeding user-defined missingness thresholds can be excluded from the analysis, while monomorphic sites can optionally be retained. Importantly, because the number of pseudohaplotypes contributing to a given window depends on how missingness is handled, filtering at the haplotype level can reduce the total number of observed haplotypes within a window. As a result, the truncated HFS used by SL may vary across windows as a function of missingness, an effect we explicitly examine in our simulations.

### Ancient DNA data

We downloaded the Allen Ancient Data Resource (Mallick *et al*., 2024) v54.1 on April 6, 2023. We excluded individuals with fewer than 25,000 SNPs, lacking geographic coordinates, without UDG treatment, or with a deamination rate at the first base of sequencing reads below 0.03. We identified individuals from two of the ancestral European populations (Anatolian Early Farmers (EF), Western hunter gatherers (WHG)) and from two admixed populations, Central European EF and Central Europe Late Neolithic/Bronze Age (LNBA). We placed individuals into our four population groups based on sample age and geographical origin (i.e. the country in which the sample was excavated). After filtering and sample selection, our dataset contained 435 individuals spanning the pre-Neolithic to the Bronze Age across Anatolia and the European continent. We performed PCA with smartpca in EIGENSOFT v7.2.1 (Price *et al*., 2006; Patterson *et al*., 2006) to confirm clustering within each group. For each resulting group, we computed the distribution of missing data and used these empirical distributions to introduce missingness into the corresponding simulated populations.

### Empirical data analysis

We applied SF2, SL, and G12 to empirical data from four ancient populations (Anatolia EF, WHG, Central Europe EF, and LNBA; **SI Table 1**) to identify putative sweep signals in unadmixed and admixed groups. Anatolia EF and WHG represent largely unadmixed ancestral populations, whereas Central Europe EF shows admixture between early farmers and Western Hunter-Gatherers, and LNBA further includes substantial Steppe-related ancestry. All methods were run using the same parameters as in the simulations. Candidate sweeps were defined as peaks whose statistic values exceeded the 1% false positive rate threshold derived from the neutral simulations. We annotated all genes overlapping each peak and defined sweep regions by merging clusters of outlier genes within 1 Mb, following a procedure similar to Souilmi et al. (2022), to account for the fact that a single sweep footprint can extend across multiple adjacent genes. Genes fully contained within the merged interval were then added to the region.

## Results

To enable application to low-coverage ancient DNA data, we implemented a modified version of saltiLASSI (SL) incorporating tolerance-based haplotype clustering and missingness filtering (see Methods), and compared its performance to SweepFinder2 (SF2) and G12, which were applied as originally described (Souilmi *et al*., 2022; Pandey *et al*., 2024).

### Performance in ancestral populations

We simulated three ancestral populations—Anatolia Early Farmers (EF), Western Hunter-Gatherers (WHG), and Steppe (see Methods; **Fig. 1**)—to evaluate the performance of saltiLASSI (SL), SweepFinder2 (SF2), and G12 under non-admixed demographic histories. Overall, SL and G12 were more powerful than SF2 for some sweep scenarios, especially at weaker selection coefficients, although power for all methods increased with both selection strength and sweep recency **(Fig. 2)**.

**Figure 1.**
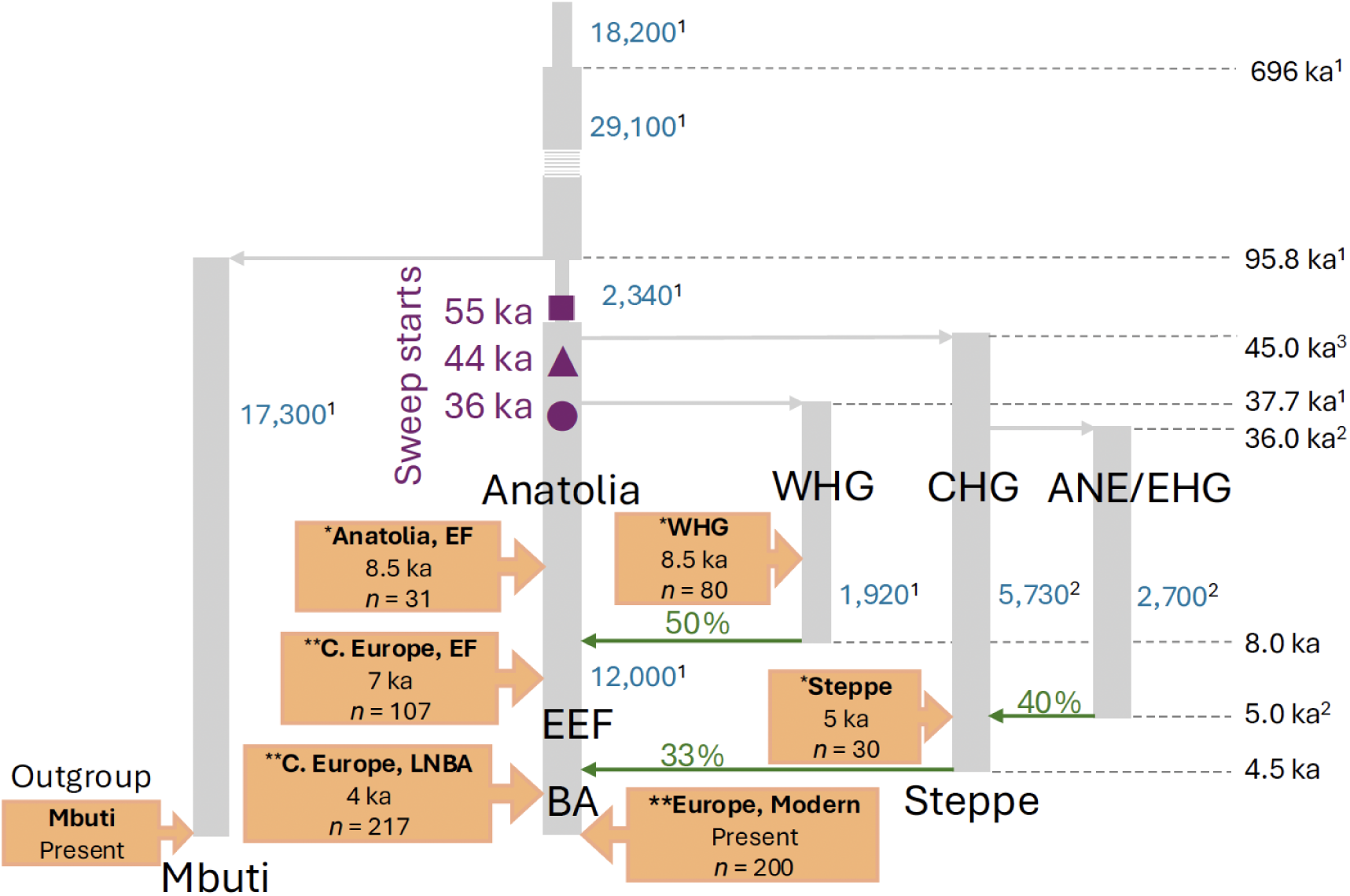
European demography model: Diagram of the European demography modeled in SLiM4. Three sweep start times (purple shapes) were simulated before and after two population splits. Orange boxes represent the populations sampled at specific time points and sample sizes. Blue numbers indicate the simulated effective population size for each population. Green arrows and numbers indicate direction and proportion of admixture. Numerical superscripts reference where certain model parameters were initially estimated (^3^Jones *et al*., 2015; ^2^Damgaard *et al*., 2018; ^1^Kamm *et al*., 2020). Figure was adapted from Souilmi et al., 2022. *ancestral population, **admixed population

**Figure 2.**
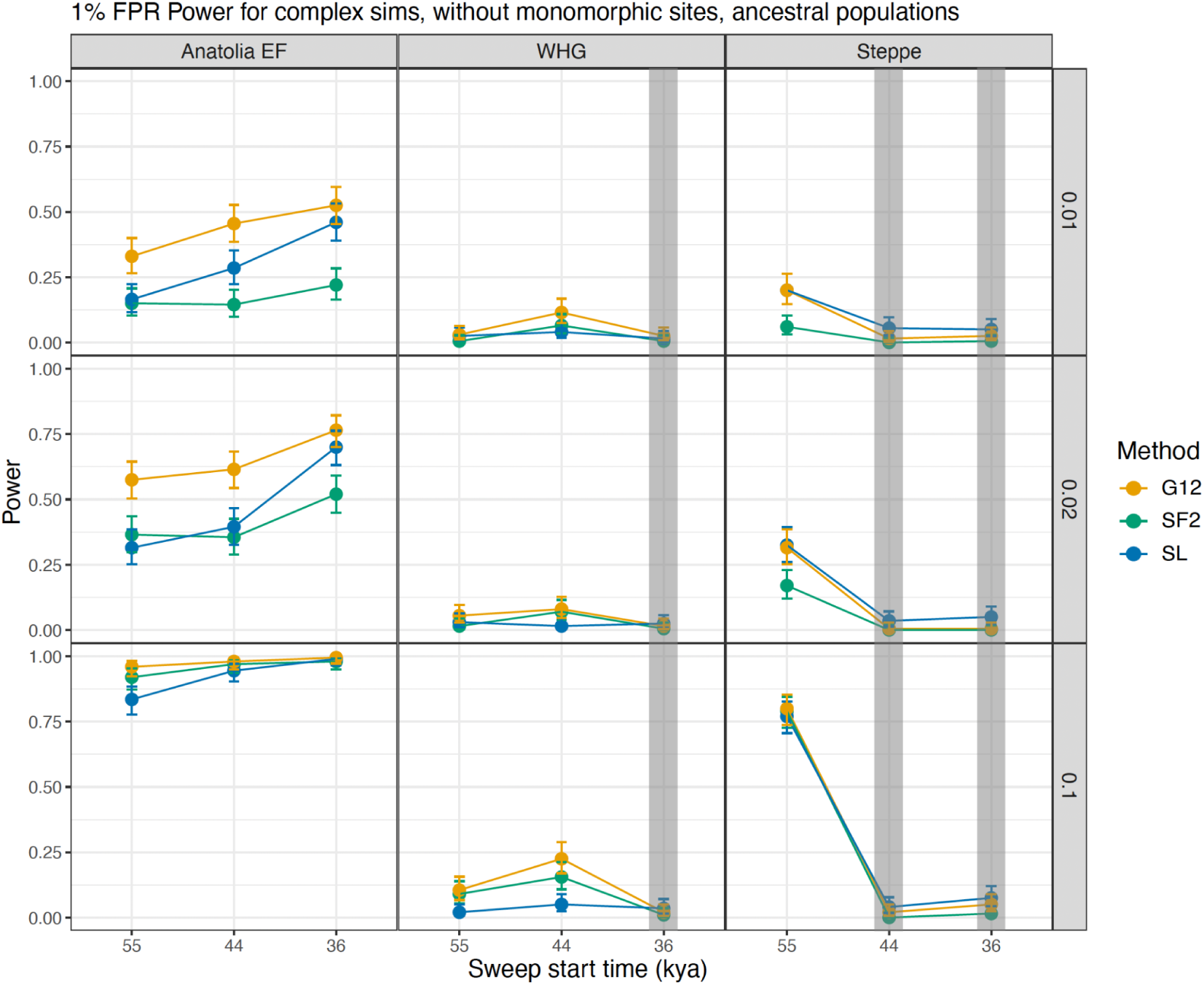
Ancestral population 1% FPR power for detecting hard sweeps: Each panel illustrates the power to detect hard sweeps for three selective sweep scan methods (G12, yellow; SF2, green; SL, blue). Power was calculated for 400 simulations run for each population (columns) under three different selection coefficients (rows; *s* = 0.01, 0.02, 0.1) and three different start times. Grey boxes indicate that the sweep did not occur in that population.

As selection strength increased, power rose markedly, particularly for sweeps beginning around 55 kya in Anatolia EF and Steppe. Under intermediate selection (*s* = 0.02), SL detected sweeps with 32% power (95% CI [0.25-0.38]; **Fig. 2**; **SI Table 2**) in Anatolia EF, increasing to 83% (95% CI [0.78-0.88]) at *s* = 0.1. SF2 displayed similar, but slightly higher, power (37% [0.30-0.44] at *s* = 0.02 and 92% [0.87-0.95] at *s* = 0.1). In contrast, G12 exhibited significantly higher power than both SL and SF2 at *s* = 0.02 (*p* < 0.0001; **Fig. 2**; **SI Table 3**) and higher power than SL at *s* = 0.1 (*p* < 0.0001), reaching 58% (95% CI [0.50-0.64]) and 96% (95% CI [0.92-0.98]), respectively.

Power to detect selective sweeps in WHG was low across all methods and selection scenarios (**Fig. 2**). In the Steppe population, SL and G12 performed similarly under low and intermediate selection strengths, whereas SF2 showed significantly lower power (*p* < 0.001 and *p* < 0.01; **Fig. 2**; **SI Table 3**). Under strong selection, however, there were no significant differences in power between methods (*p* > 0.5; **SI Table 3**) and power increased to an average of 84%. Overall, these results indicate that haplotype-based statistics perform better than SFS-based statistics for detecting relatively recent sweeps in non-admixed populations, especially at lower selection strengths, whereas all methods perform similarly when selection is strong.

### Performance in admixed populations

We next consider admixed populations, where the detection of selective sweeps is complicated by ancestry mixing and the potential dilution of sweep signals over time. We distinguish between *continuous selection*, in which selection persists after admixture, and *non-continuous selection*, in which selection ceases at the time of admixture and sweep signals remain diluted in subsequent generations.

As in the ancestral populations, power to detect sweeps increased with selection strength, but patterns in admixed populations were more complex (**Fig. 3**). We first consider continuous selection. Under very weak selection (*s* = 0.01), power was low across all three methods and populations, with none of the methods consistently detecting sweeps, reflecting limited post-admixture increase of the beneficial allele frequency (**Fig. 3)**. At intermediate selection strength (*s* = 0.02), differences between methods became apparent. SL exhibited modest but relatively stable power across sweep ages, whereas G12 and SF2 showed pronounced declines as sweep signals were diluted by admixture. This contrast was most evident in the highly admixed Central Europe LNBA population, where SL achieved significantly higher power (34%, 95% CI [0.27-0.40], *p* < 0.0001; **Fig. 3**; **SI Table 4-5**) than both G12 and SF2 for the most recently introduced sweep. These results suggest that SL is less sensitive to admixture-driven dilution of sweep haplotypes and retains power to detect partially eroded sweep signals.

**Figure 3.**
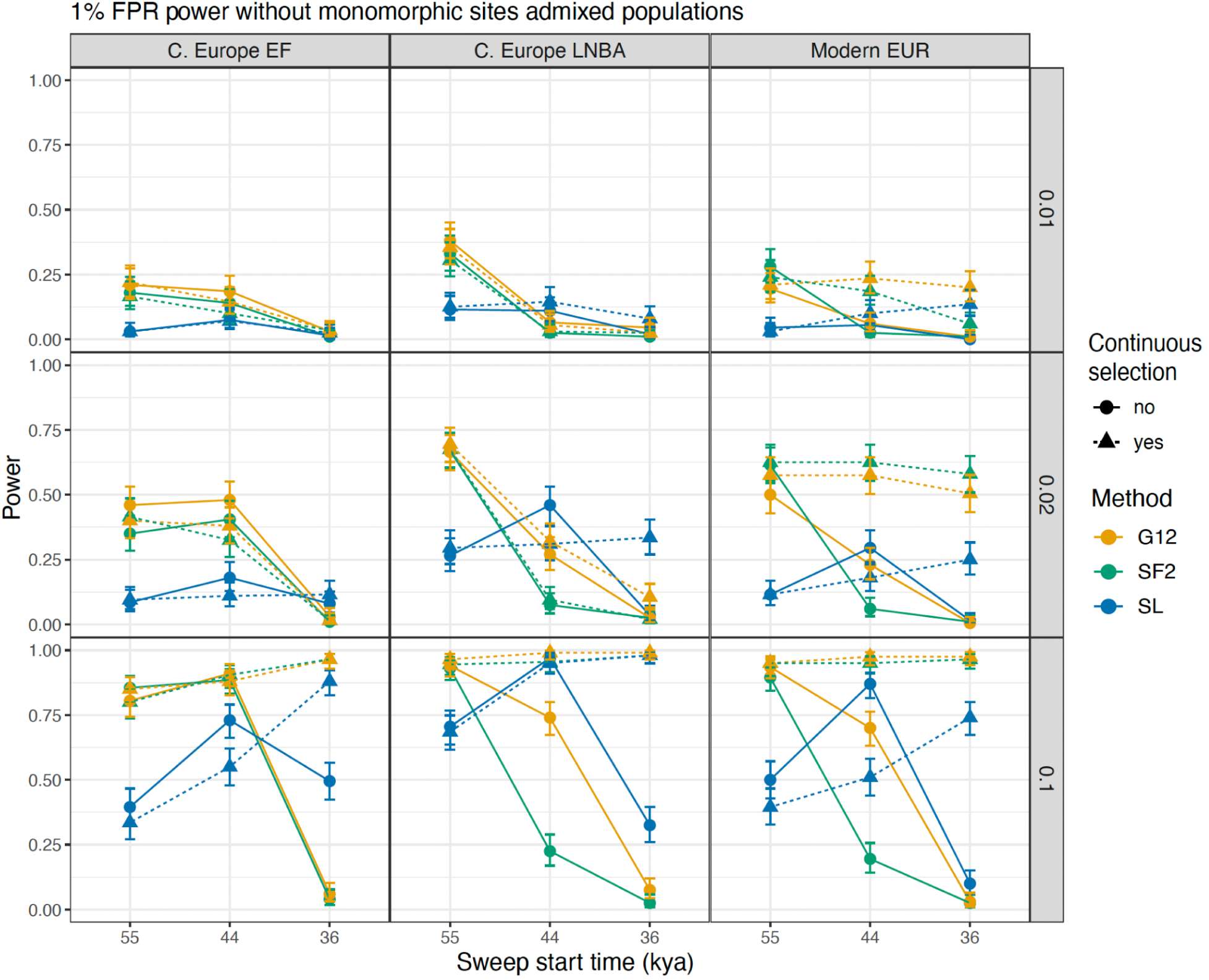
Admixed populations 1% FPR power for detecting hard sweeps: Each panel illustrates the power to detect hard sweeps for three selective sweep scan methods (G12, yellow; SF2, green; SL, blue). Power was calculated for 200 simulations run for each population (columns) under three different selection coefficients (rows; *s* = 0.01, 0.02, 0.1), continuous (triangle, dashed lines) and non-continuous (circle, solid lines) selection, and three different start times.

Under strong continuous selection (*s* = 0.1), power increased substantially for all methods. In this case, selection rapidly elevated the frequency of the beneficial allele after admixture, effectively re-establishing a coherent sweep signal. While G12 and SF2 achieved the highest absolute power (remaining above 80%), SL again showed competitive performance in Central Europe LNBA for recent sweeps (98%, 95% CI [0.95-0.99]), consistent with enhanced robustness to admixture when selection remains strong (**Fig. 3**).

Under non-continuous selection, differences in method performance became more pronounced. In Central European EF, SL showed lower power than G12 and SF2 for older sweeps when *s* ≥ 0.02. However, for recent sweeps under strong selection (*s* = 0.1), SL significantly outperformed both methods (*p* < 0.0001; **Fig. 3**; **SI Table 5**). Interestingly, SL achieved the highest power of the three methods for sweeps starting ∼44 kya in Central European LNBA and modern Europeans at *s* ≥ 0.02, although its advantage over G12 was not significant in modern Europeans at *s* = 0.02 (*p* > 0.2; **Fig. 3**; **SI Table 5)**. As sweep signals became increasingly diluted, comparing sweeps beginning ∼44 kya to those beginning ∼36 kya (which experience more admixture), SL’s power under strong selection (*s* = 0.1) declined substantially, from 97% (95% CI [0.94–0.99]) to 32.5% (95% CI [0.26–0.39]) in Central European LNBA and from 87% (95% CI [0.82–0.91]) to 10% (95% CI [0.06–0.15]) in modern Europeans (**Fig. 3**; **SI Table 4**).

G12 and SF2 showed no significant differences in power for older sweeps (*p* > 0.05; **Fig. 3**; **SI Table 5**). However, for sweeps starting ∼44 kya and *s* ≥ 0.02, G12 significantly outperformed SF2 in both Central European LNBA and modern Europeans (*p* < 0.0001). For recent sweeps, both methods showed low power (<10%) across all admixed populations, regardless of selection strengths (**Fig. 3**; **SI Table 4**). Overall, non-continuous selection combined with increasing admixture produced a distinct pattern of power in SL that was not observed for G12 or SF2.

However, no single method consistently outperformed the others across all scenarios, suggesting that combining multiple approaches may provide a more comprehensive set of candidate sweep regions particularly in admixed populations.

### Effect of including monomorphic sites

It is standard practice to filter out monomorphic sites prior to analysis. However given the limited amount of data and high levels of missingness in ancient DNA, removing invariant sites may discard informative signal. To evaluate this, we applied haplotype-based methods to simulated data both before and after filtering monomorphic sites. We did not apply SF2 to data excluding invariant sites, as its performance in ancient populations has been previously established (Souilmi *et al*., 2022), and here it serves primarily as a benchmark for haplotype-based approaches.

Retaining invariant sites increased SL’s power overall, particularly for sweeps under continuous selection and for older sweeps (∼44 kyo and earlier) under non-continuous selection (**Fig. 4**; **SI Table 6).** Power gains were most pronounced at higher selection strengths and the oldest sweeps. In Anatolia EF, SL’s power to detect recent sweeps increased from 9.5% (95% CI [0.06-0.14]; **SI Table 4**) to 20% (95% CI [0.15-0.26]; **SI Table 6**) under weak selection, from 18% (95% CI [0.13-0.24]; **SI Table 4**) to 48% (95% CI [0.40-0.55]; **SI Table 6**) under intermediate selection, and from 53% (95% CI [0.46-0.60]; **SI Table 4**) to 93% (95% CI [0.88-0.96]; **SI Table 6**) under strong selection.

**Figure 4.**
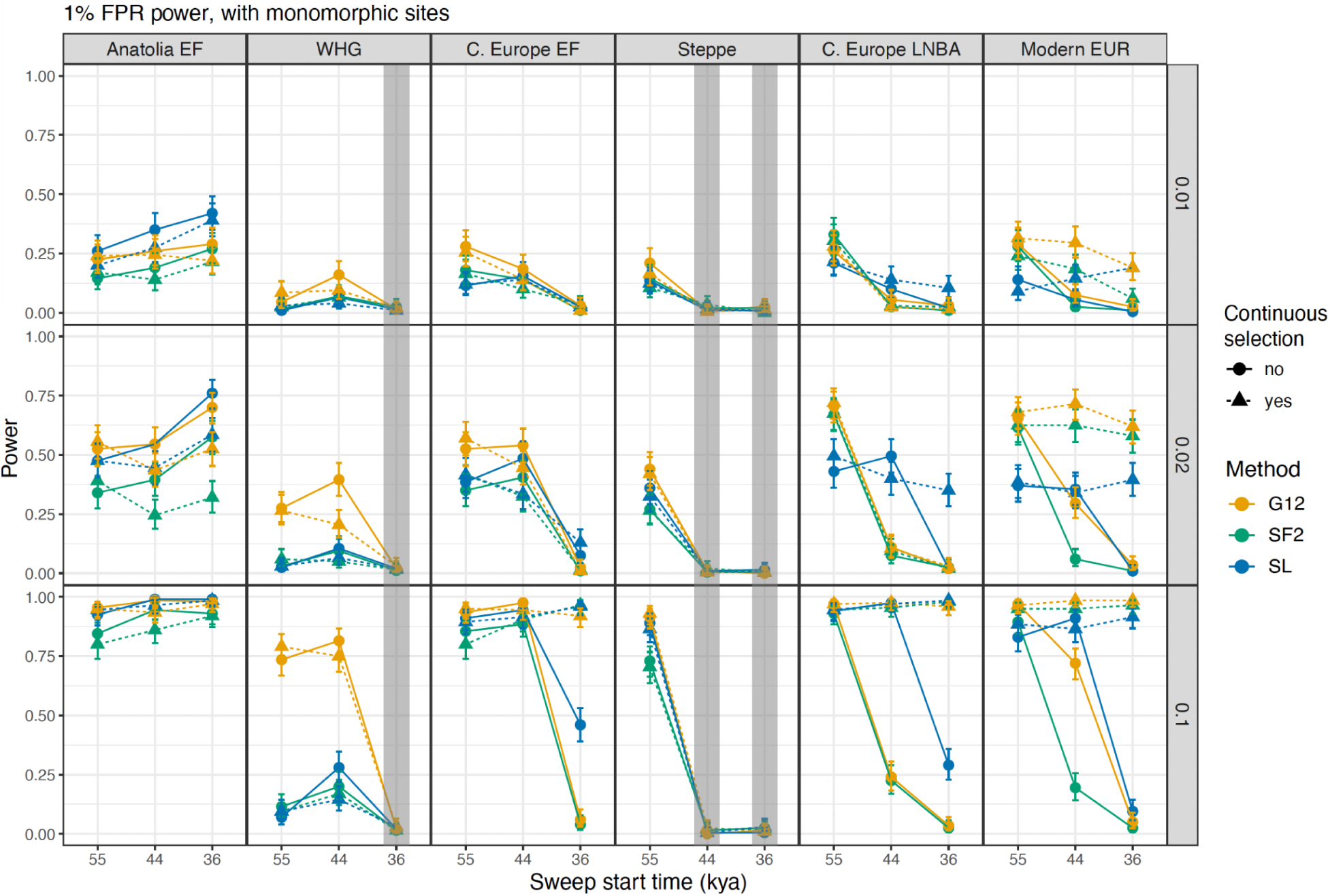
All populations 1% FPR power for detecting hard sweeps while retaining invariant sites: Each panel illustrates the power to detect hard sweeps for three selective sweep scan methods (G12, yellow; SF2, green; SL, blue). Power was calculated for 200 simulations run for each population (columns) under three different selection coefficients (rows; *s* = 0.01, 0.02, 0.1), continuous (triangle, dashed lines) and non-continuous (circle, solid lines) selection, and three different start times. Grey boxes indicate that the sweep did not occur in that population.

Including monomorphic sites had the greatest impact on power in Central Europe EF compared to the other admixed populations. When keeping monomorphic sites, power to detect sweeps 55kyo in Central Europe EF when *s* = 0.02 was about 9% for both continuous and non-continuous selection **(Fig. 3**; **SI Table 4)**. Retaining monomorphic sites increased power by ∼30–32% under these conditions and by over 50% under strong selection (*s* = 0.1; **Fig. 4**; **SI Table 6**). Modern Europeans had a similar increase in power for both selection schemes for our oldest sweeps (about 26% when *s* = 0.02 and 33-49% when *s* = 0.1), but not Central European LNBA. However, power marginally improved for recent sweeps in all admixed populations under non-continuous selection (**Fig. 4**; **SI Table 6**).

G12 also showed increased power when monomorphic sites were retained, particularly for older sweeps under both continuous and non-continuous selection (**Fig. 4**; **SI Table 6**). The largest improvement was observed in WHG, where power increased from below 23% (**Fig. 2**; **SI Table 2**) to over 73% at *s* = 0.1 (**Fig. 4**; **SI Table 6**). However, G12’s power decreased in Central Europe LNBA under non-continuous selection for sweeps beginning 44 kya when *s* ≥ 0.02. We address why this might be happening below.

Overall, retaining invariant sites substantially improved sweep detection, particularly for SL, which performed as well as or better than SF2 and G12 for recent sweeps in admixed populations under strong, non-continuous selection (**Fig. 4**; **SI Table 7**). These results suggest that including monomorphic sites can enhance power in low-coverage ancient DNA data and improve the robustness of haplotype-based methods across a range of demographic and selective scenarios.

### Effect of window size

We evaluated two different window sizes (51 and 101 SNPs); results presented above correspond to 51 SNP windows. Here, we compare power across window sizes for SL and G12.

Window size had a strong effect on SL, but only a limited effect on G12 (**Fig. 5**; **SI Figs. 1–3**). For SL, smaller windows (51 SNPs) generally yielded higher power than larger windows, with the largest differences observed in admixed populations when comparing between the same sweep conditions (**Fig. 5**; **SI Fig. 3; SI Tables 4, 6, 8-9**). However, larger windows significantly improved SL’s performance for recent sweeps in ancient admixed populations under strong, non-continuous selection (*p* < 0.01; **SI Tables 10–11**). Retaining monomorphic sites further enhanced SL’s power under these conditions (**SI Fig. 3; SI Tables 6, 9**). In Central Europe EF and LNBA, larger window sizes power increased by 62% and 133%, respectively, when including monomorphic sites, compared to increasing by 28% and 94% when excluding monomorphic sites (**Fig. 5**; **SI Fig. 3; SI Tables 4, 6, 8-9**).

**Figure 5.**
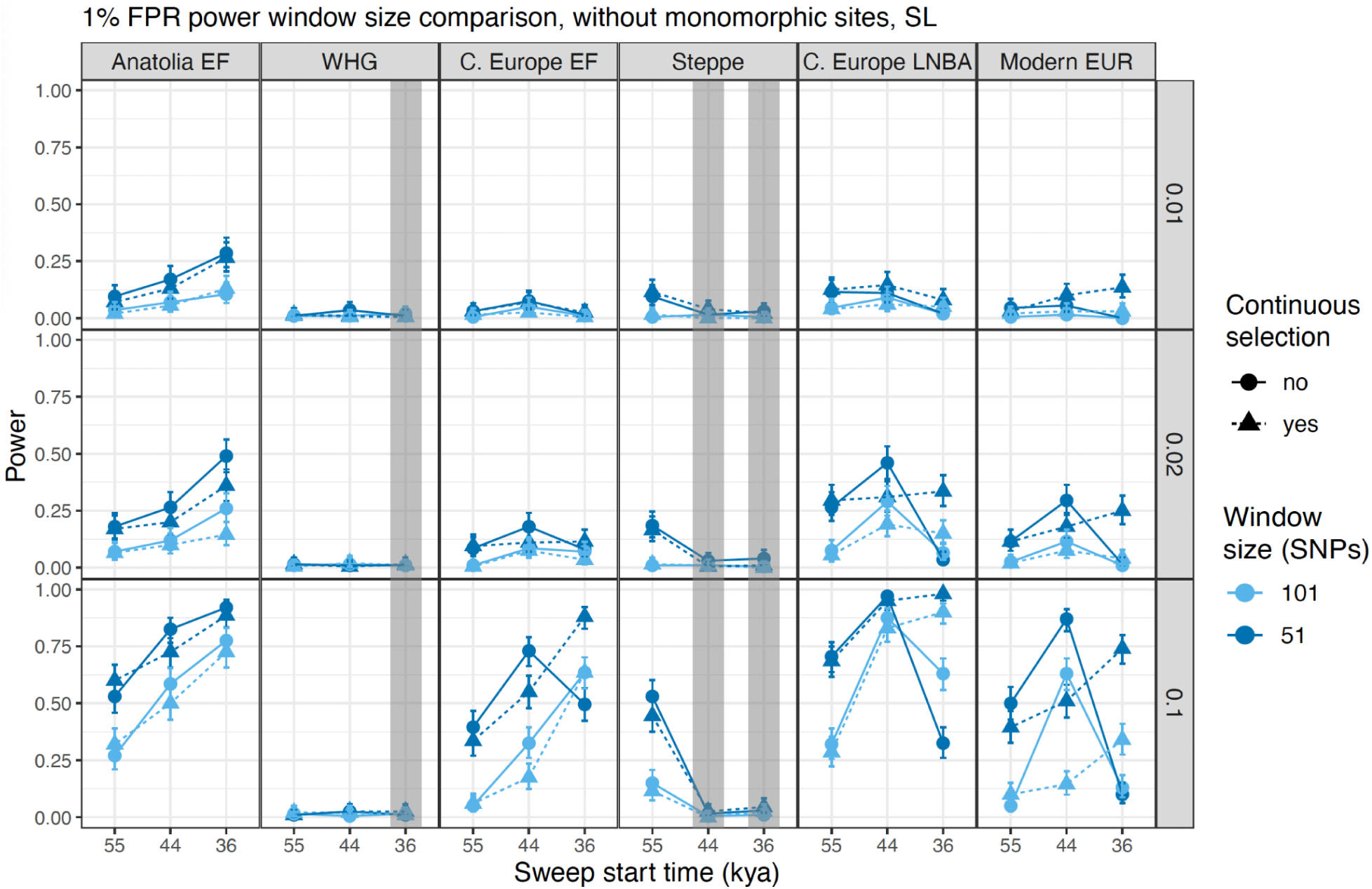
Window size impacts power to detect hard sweeps: Each panel illustrates SL’s power to detect hard sweeps for two different window sizes: 51 (dark blue) and 101 (light blue) SNPs. Power was calculated for 200 simulations run for each population (columns) under three different selection coefficients (rows; *s* = 0.01, 0.02, 0.1), continuous (triangle, dashed lines) and non-continuous (circle, solid lines) selection, and three different start times. Grey boxes indicate that the sweep did not occur in that population.

In contrast, window size had relatively little impact on G12 overall (**SI Figs. 1–2**). Larger windows modestly increased power in highly admixed populations for sweeps beginning 44 kya when *s* ≥ 0.02 under non-continuous selection, both when including and excluding monomorphic sites (*p* < 0.0001**; SI Fig. 1; SI Tables 10–11**). Larger window sizes might be more accurately capturing patterns of the sweeping haplotype that have not been fully broken up by admixture whereas smaller window sizes might give too fine of resolution to detect historic sweep patterns, thus power increases with larger windows. However, a few exceptions were observed: smaller windows significantly increased power in WHG for data with invariant sites (*p* < 0.01), as well as in modern Europeans for older sweeps under weak and intermediate selection (*p* < 0.03; **SI Fig. 2; SI Table 11**).

Overall, SL performs best with smaller windows, reflecting its sensitivity to local haplotype structure, whereas G12 is relatively insensitive to window size and performs slightly better with larger windows. Under strong admixture and recent sweeps, however, larger windows improve SL’s performance, indicating that broader windows help recover diluted sweep signals.

### Sweep detection in empirical data

We applied SL, SF2, and G12 to four empirical populations (**SI Fig. 4**): Anatolia EF (*n* = 31), WHG (*n* = 80), Central Europe EF (*n* = 107) and Central Europe LNBA (*n* = 217), using the same parameter settings as in the simulations. Consistent with the simulation results, window size and the inclusion of monomorphic sites had a strong impact sweep detection with SL. Smaller windows (51 SNPs) identified more sweeps than larger windows (101 SNPs) (25 vs. 19 sweeps). Including monomorphic sites substantially increased the number of detected sweeps for both window sizes, to 123 and 83 sweeps for smaller and larger windows, respectively **(Fig. 6A-D**; **SI Tables 12-13)**. Specifically, 96% (24/25) of sweeps identified with smaller windows and 84% (16/19) with larger windows were recovered when monomorphic sites were included. Across all populations, we identified 32 sweeps when excluding monomorphic sites and 144 when including them (**Fig. 7A**), with 94% (30/32) of sweeps identified when excluding monomorphic sites also present in the inclusive dataset. SF2 identified 20 sweeps **(SI Table 14)**.

**Figure 6.**
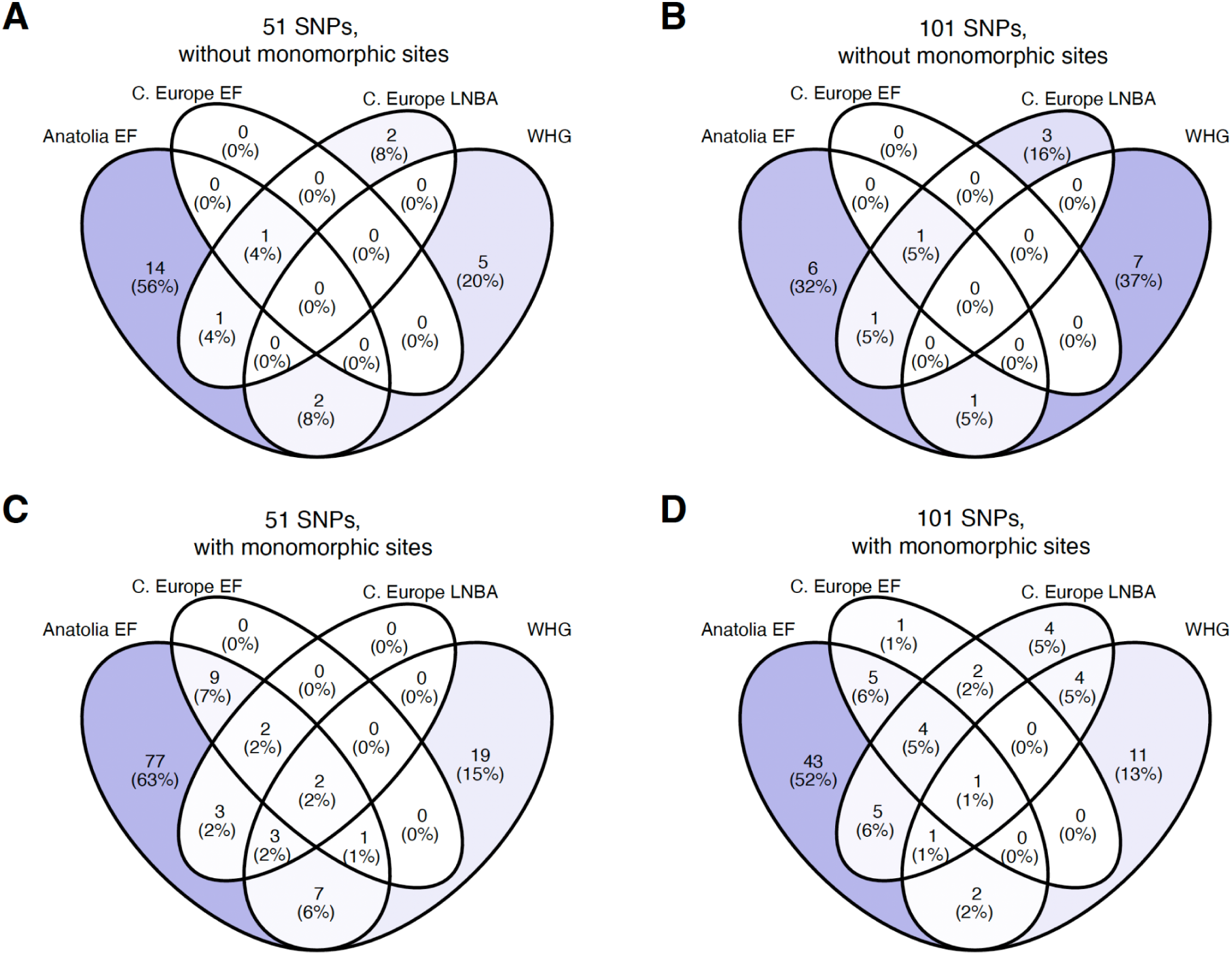
Venn diagrams of SL sweeps shared between populations: Panels A-D show the number of sweeps identified via SL that are shared between four ancient populations (WHG, Anatolia EF, central European EF, and central European Bronze Age). The number of sweeps identified are based on two major method parameters: window size and filtering of monomorphic sites. We tested two window sizes (101 and 51 SNPs).

**Figure 7.**
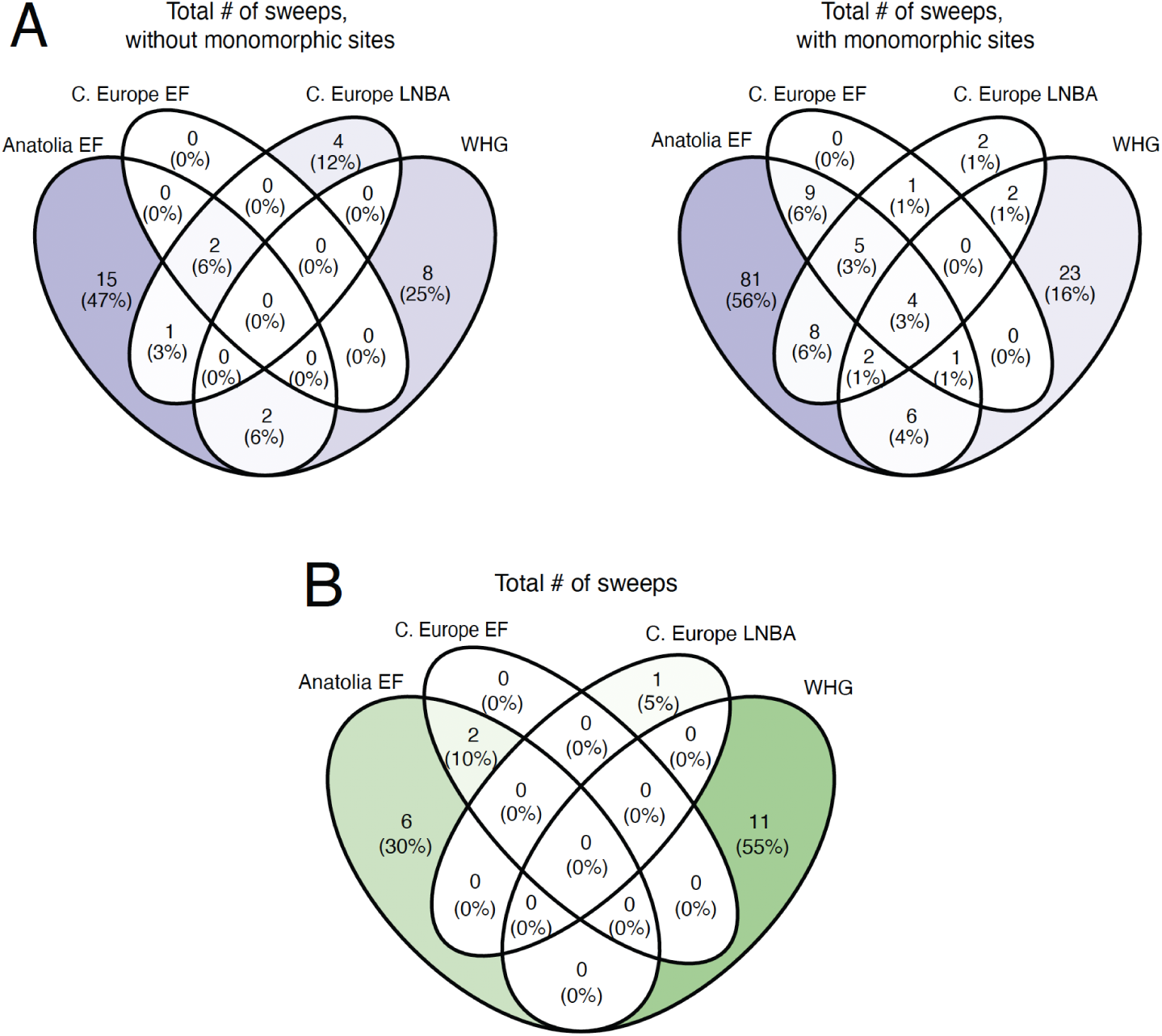
Venn diagrams of overlapping sweep regions between ancient populations: A-B compare the number of unique and overlapping sweeps for each population identified by SL (A), and SF2 (B). A: Sweeps identified by both window sizes were combined into a single sweep region.

G12 showed markedly different behavior from SL, identifying substantially more sweep regions than either SL or SF2. As with SL, smaller windows (51 SNPs) identified more sweeps than larger windows (814 vs. 435 sweeps; **SI Fig. 5A–B; SI Table 15**). Including monomorphic sites further increased the number of detected sweeps for smaller windows (919 sweeps) but slightly decreased the number for larger windows (429 sweeps; **SI Fig. 5C–D; SI Table 16**).

Moreover, G12 identified substantially larger sweep regions than either SL or SF2. In the dataset excluding monomorphic sites, the average sweep length was ∼4.5 Mb for 101 SNP windows and∼0.6 Mb for 51 SNP windows (**SI Table 15**). Similar patterns were observed when including monomorphic sites (∼4.6 Mb and ∼0.8 Mb; **SI Table 16**). In contrast, sweep lengths for SL and SF2 were typically below 1 Mb, with a maximum of 2.3 Mb for SL (**SI Tables 12–14**), consistent with previous studies reporting sweep lengths generally below ∼4 Mb and typically under 1 Mb (Souilmi et al. 2022; Pandey et al. 2024).

Given the unexpectedly large size and number of sweeps detected by G12, we performed a sensitivity analysis using a more conservative neutral cutoff, obtained from simulations with a recombination rate set to half the original value (*r* = 5 x 10^-9^). This higher threshold reduced sweep lengths substantially, with most sweeps falling below 1 Mb, although some remained as large as 7.4 Mb. However, this also led to an increase in the total number of detected sweeps, particularly for the larger window size (**SI Tables 17–18**). In contrast, the same adjustment reduced the number of detected sweeps for both SL and SF2 (**SI Tables 19–21**).

Overall, G12 produced an unusually large number of broad sweep regions that were highly sensitive to parameter choices and inconsistent with expectations from both simulations and previous studies. These results suggest that G12 should be applied with caution in low-coverage or highly admixed datasets, where it may overestimate the extent and number of sweep signals, particularly in the presence of recombination-rate heterogeneity. Accordingly, careful calibration of thresholds and validation against simulations are essential when applying G12 in such contexts; given these issues, we did not pursue further analyses of G12-detected sweeps.

### Sweeps shared between populations

Next, we investigated the extent to which sweeps are shared between populations. Overall, SL identified more shared sweeps than SF2, although the number of shared signals depended strongly on whether monomorphic sites were included.

When excluding monomorphic sites, SL did not detect any sweeps shared across all four populations. In contrast, when including monomorphic sites, SL identified four sweeps shared among all populations. Expanding the analysis to sweeps shared between at least two populations, 16% (5/32) and 24% (34/144) of sweeps were shared when excluding and including monomorphic sites, respectively (**Fig. 7A**). Of these, more than half were shared between at least one ancestral and one admixed population (**Fig. 7A; SI Tables 12–13**). In comparison, SF2 identified only two shared sweeps, both between Anatolia EF and Central Europe EF (**Fig. 7B**). Using a more stringent neutral cutoff, calculated from simulations with a lower recombination rate (r = 5 × 10⁻⁹), reduced the number of shared sweeps detected by both SL and SF2. For SF2, the two shared sweeps were reduced to a single shared signal (**SI Fig. 6**). For SL, only a small number of shared sweeps remained, none of which were shared across all four populations: one sweep was shared among Anatolia EF, Central Europe EF, and LNBA; one between Anatolia EF and Central Europe LNBA; and three between Anatolia EF and WHG (**SI Fig. 7A–D**).

Including monomorphic sites and using larger window sizes modestly increased the number of shared sweeps detected by SL (**SI Fig. 7A–D**). Overall, these results indicate that parameter choices, particularly the inclusion of monomorphic sites and the definition of neutral thresholds, can influence the detection of shared sweeps across ancient populations, and that reported patterns of sweep sharing should be interpreted in light of these sensitivities.

### Overlap with previous studies and functional relevance

To assess concordance between methods, we compared sweeps identified by SL and SF2 using datasets including monomorphic sites. Overall, 75% (14/20) of SF2 sweeps overlapped with SL sweeps, although these represented only ∼10% of SL-identified sweeps (**SI Table 22**). Most shared sweeps were detected in the same populations, but one sweep was detected by SF2 in WHG and by SL in Anatolia EF, indicating that the two methods can capture partially distinct signals.

Consistent with our simulation results, SL identified substantially more sweeps in admixed populations than SF2. In Central Europe EF and LNBA, SL detected 34 sweeps compared to only three for SF2, two of which overlapped with SL (**Fig. 7A–B**; **SI Table 22**).

We next compared our results to previously reported sweep regions in ancient populations (Souilmi *et al*., 2022) and modern populations (Racimo, 2016). Among SL sweeps, 32 of 144 overlapped with regions reported by Souilmi et al. (2022), including 12 sweeps detected in populations not previously reported in those regions (**SI Table 23**). In contrast, 15 of 20 SF2 sweeps overlapped with Souilmi sweeps and were found in the same populations (**SI Table 24**).

Comparison with sweeps identified in modern populations revealed that 19 of 144 SL sweeps and 4 of 20 SF2 sweeps overlapped with regions reported by Racimo (2016) across European, East Asian, and ancestral Eurasian populations (**SI Tables 25–26**). Only two sweeps were detected by both methods in these regions. Notably, a majority of SL overlaps (∼53%) were specific to modern European populations, with smaller proportions unique to East Asian and ancestral Eurasian groups.

Finally, enrichment analysis using FUMA (Watanabe et al. 2017) revealed that sweeps identified by both methods were enriched for GWAS categories related to autoimmune diseases, neurological traits, physical traits, and immunity (**SI Fig. 8**; **SI Table 27**). However, SL uniquely identified additional enriched categories, including height, malaria, and hepatitis B–related traits. In the HLA region, SL identified all genes detected by SF2, including previously reported targets such as ZKSCAN3 *(Mathieson et al., 2015)*, but also detected two sweep regions that include multiple olfactory receptor (OR) genes. Interestingly, these OR genes were associated with anthropometric traits such as hip circumference (Christakoudi *et al*., 2021) and height (Tachmazidou *et al*., 2017; Schoeler *et al*., 2025), but also psychiatric traits (Zhao *et al*., 2021) and diet (May-Wilson *et al*., 2022). Furthermore, OR genes have been linked to sex-based differences (Förster *et al*., 2025) and obesity and BMI (Ramos-Lopez *et al*., 2019).

Overall, these results demonstrate substantial concordance with previous studies and highlight the biological relevance of the detected sweep regions, while also indicating that SL captures additional candidate sweeps, particularly in admixed populations.

## Discussion

Historical data provides unique opportunities to study how selection has shaped modern populations, but signals of selection can be obscured by demographic processes such as admixture and drift (Pandey *et al*., 2024). Admixture is pervasive in human history and can result in masking historical selection signals (Souilmi *et al*., 2022; Pandey *et al*., 2024). Here, we developed and evaluated a haplotype-based approach tailored to low-coverage ancient DNA data and applied it to both simulated and empirical datasets to access its performance relative to existing methods.

### Ideal selection scan parameters

Our results show that there is no single optimal approach for detecting selective sweeps across all demographic scenarios. Instead, combining complementary methods and carefully tuning parameter choices is likely to yield the most reliable results, particularly when demographic history is complex or unknown.

In particular, the inclusion of monomorphic sites substantially improves the performance of haplotype-based methods in low-coverage ancient DNA. While monomorphic sites are typically removed prior to analysis, doing so in aDNA may discard informative signal and reduce power. Consistent with this, retaining invariant sites increased power for both SL and G12, although the magnitude of this effect varied across populations and selection regimes. For SL, including monomorphic sites was critical for achieving comparable or improved performance relative to SF2, particularly in admixed populations and for non-continuous selection scenarios.

Power remained low in populations with long-term small effective population size (e.g. WHG), even under strong selection, reflecting the difficulty of detecting sweeps in highly homozygous backgrounds. In contrast, G12 showed greater robustness under these conditions, suggesting that different methods capture complementary aspects of haplotype structure. The reduced performance of SL in such settings may reflect its use of a likelihood framework based on the full HFS, which may be less sensitive when haplotype diversity is limited. Notably, SL was not originally designed to incorporate monomorphic sites, indicating that further methodological refinement may improve its performance in low-diversity contexts.

Window size also had a strong impact on detection power. Smaller windows improved sensitivity to older sweeps by capturing localized haplotype structure, whereas larger windows enhanced detection of recent sweeps in admixed populations. This pattern is consistent with previous work showing that larger window sizes allow for better detection of recent and soft sweeps (DeGiorgio and Szpiech, 2022). Thus, this likely reflects the breakdown of sweeping haplotypes following admixture, which can transform hard sweeps into softer, more diffuse signals (Pennings and Hermisson, 2006). Larger windows may partially recover these signals by integrating information across fragmented haplotypes. In contrast, G12 was comparatively insensitive to window size, with performance more strongly influenced by admixture and the continuity of selection. While admixture may reduce the frequency of the most common haplotype, source populations with low diversity contribute fewer novel haplotypes, minimizing shifts in the haplotype frequency spectrum particularly in sweep regions where the HFS is already distorted (Lohmueller *et al*., 2010). This distinction is important because SL leverages the full HFS while G12 focuses on the relative frequencies of the most common haplotype for determining sweep regions. Taken together, these results demonstrate that parameter choices—particularly the inclusion of monomorphic sites and the choice of window size—are critical for optimizing sweep detection in ancient DNA. In practice, we recommend retaining invariant sites and exploring multiple window sizes when applying haplotype-based methods, especially in admixed or low-coverage datasets.

### Method performance in empirical data

Previous studies have shown that SF2 and G12 have limited power to detect selective sweeps in admixed populations (Souilmi *et al*., 2022; Pandey *et al*., 2024). More recently, Harris et al. (2026) applied a domain-adapted neural network (DANN) that included a gradient reversal layer (GRL) and successfully detected sweeps across time in ancient European populations and moderately improved sweep detection power, even with model mis-specification (Harris *et al*., 2026). While this approach ameliorates, to some degree, the dependence on an accurately specified demographic model, mismatch still remains a problem (Mo and Siepel, 2023).

Moreover, these approaches require extensive training data, limiting their applicability to systems where demographic history is uncertain or poorly characterized (Harris, Sackman, *et al*., 2018).

Pandey et al. (2024) and Harris et al. (2026) demonstrated that G12 successfully detected well known sweep regions in ancient data, however while our G12 results showed high power in simulations, its empirical behavior was inconsistent, producing unrealistically large numbers of sweep regions and strong sensitivity to recombination rate and window size. This discrepancy could be for a few reasons. First, both studies use a fixed window of 201 SNPs to calculate power and thresholds, whereas we use smaller, sliding windows. Second, we calculate the neutral G12 threshold for each simulated population and they calculate a single cutoff for all populations. The single cutoff leads to a larger threshold that could produce more reasonable results, and we find this to be the case in our data even with smaller window sizes. However, larger window sizes reduce the cutoff and it becomes less reasonable for all populations except C. Europe LNBA. This could be because admixture reduces overall G12 values meaning that the single cutoff is more robust to higher levels of admixture since it is calculated from multiple populations regardless of their specific demographic history. Third, our results highlight recombination rate as an additional key factor influencing sweep detection. In contrast, sweep counts from SL and SF2 decreased more predictably under changes in recombination rate (**SI Fig. 9-10**) and statistical thresholds for specific populations do not result in an unreasonable number of sweeps, suggesting greater robustness to parameter variation.

Given that G12 identified sweep regions spanning large portions of the genome, we consider these results unlikely to reflect realistic selection patterns. We therefore focus subsequent analyses on SL and SF2, which provide more consistent and interpretable signals of selection in empirical data.

### Overlap with known sweeps and functional enrichment

Consistent with our simulation results, admixture reduced the power to detect selective sweeps, with fewer sweeps identified in admixed populations than in ancestral populations. Nevertheless, SL recovered substantially more sweep signals than SF2 overall, particularly in admixed populations. Patterns observed in the empirical data closely matched those from simulations.

Using smaller window sizes with SL resulted in greater overlap with previously reported sweeps from Souilmi et al. (2022), particularly for older sweeps (**SI Table 23**), whereas larger window sizes increased the number of detected sweeps in admixed populations (**Fig. 6**). Including monomorphic sites increased the total number of detected sweeps, although this also reduced overlap with previously reported regions due to the larger number of candidate sweeps.

While most of our SF2 sweeps overlapped with sweeps reported in Souilmi et al. (2022), SF2 failed to detect several well-established sweep signals that were identified by SL. For example, the *OCA2/HERC2* region, a well known target of selection in ancient and modern European populations (Voight *et al*., 2006; Wilde *et al*., 2014; Mathieson *et al*., 2015), was detected by SL under both window sizes, but not by SF2. Notably, alleles in this region are known to be at high frequency in WHG and at intermediate frequency in Anatolia EF (Mathieson *et al*., 2015; Ju and Mathieson, 2021), suggesting that SL may be better able to capture partial or population-specific sweeps that are less detectable by SFS-based methods.

Functional enrichment analyses further supported the biological relevance of detected sweeps. Both SL and SF2 showed enrichment for GWAS categories related to neurological traits, immunity, and human physical characteristics. Notably, SL identified unique enriched categories, including height, malaria, and hepatitis B–related traits, reinforcing the value of combining complementary methods. Moreover, sweeps harboring OR genes associated with anthropometric traits (Tachmazidou *et al*., 2017; Christakoudi *et al*., 2021; Schoeler *et al*., 2025) across Anatolia EF and Central European populations might suggest additional selective pressures, potentially related to shifts in diet or environment (Ma *et al*., 2026). Together, these results demonstrate that SL not only recovers known sweep signals but also identifies additional candidate regions with plausible biological relevance, particularly in admixed populations. Combining SL with complementary approaches such as SF2 provides a more comprehensive view of selection in ancient populations.

### Conclusions

Our results highlight the central role of demography in shaping the detectability of selective sweeps and underscore the challenges of identifying selection in admixed populations. We show that combining complementary approaches such as SL and SF2 provides a more robust and comprehensive view of selection, capturing both shared signals and method-specific features of sweep regions. Importantly, parameter choices—particularly the inclusion of monomorphic sites and the choice of window size—substantially influence inference and should be carefully considered in analyses of low-coverage ancient DNA. Together, these findings demonstrate that haplotype-based methods, when appropriately parameterized, can improve the detection of selective sweeps in ancient and admixed populations and provide practical guidance for selection scans in complex demographic settings.

## Acknowledgements

We thank all the anonymous peer reviewers for productive comments on the manuscript. This work was supported by the National Institute of General Medical Sciences of the National Institutes of Health award number R35GM146926 (ZAS) and R35GM146886 (CDH). The content is solely the responsibility of the authors and does not necessarily represent the official views of the National Institutes of Health.

## Author Contributions

CDH conceived of and supervised the project. ANS performed all the simulations and data analysis. ZAS oversaw updates to the saltiLASSI software. ANS wrote the first draft of the manuscript. All authors contributed to manuscript revisions.

## Data archiving

Code to reproduce simulations can be found on GitHub at https://github.com/ans6160/haplotype-based_models_improve_sweep_detection.

