## Supplemental Figures 1-10 for "Haplotype-Based Models Improve Sweep Detection in Ancient Populations with Complex Demography"

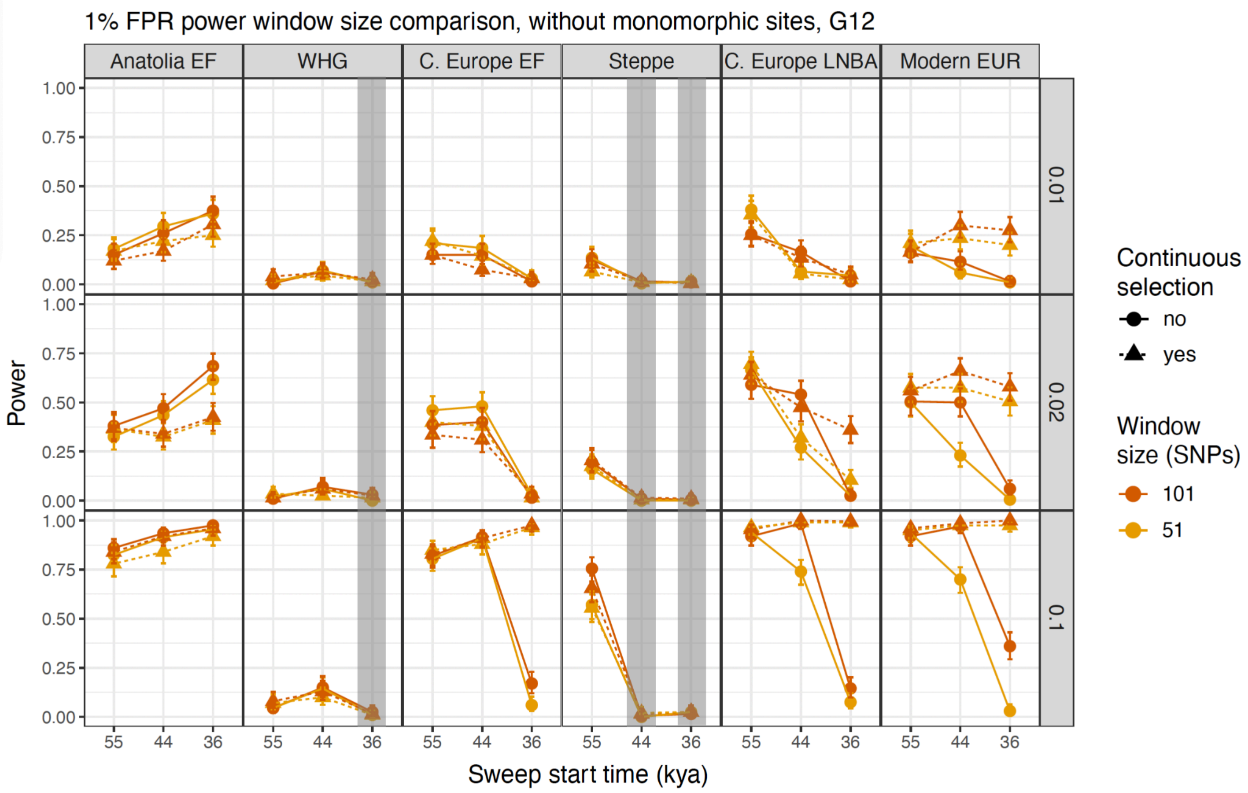


**SI Figure 1 Window size does not have a large impact on G12's power to detect hard sweeps:** Each panel illustrates G12's power to detect hard sweeps for two different window sizes: 51 (orange) and 101 (gold) SNPs. Power was calculated for 200 simulations run for each population (columns) under three different selection coefficients (rows; $s$ = 0.01, 0.02, 0.1), continuous (triangle, dashed lines) and non-continuous (circle, solid lines) selection, and three different start times. Grey boxes indicate that the sweep did not occur in that population.

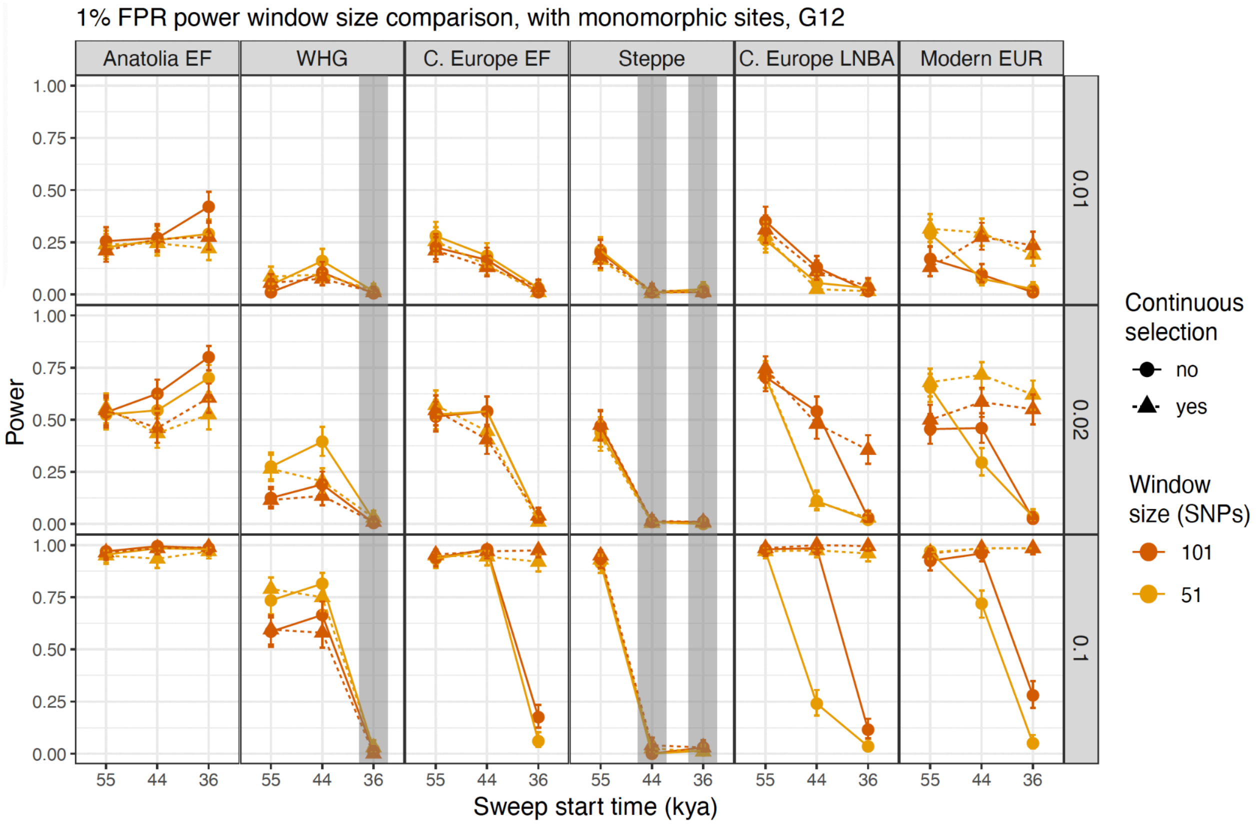


**SI Figure 2 Including monomorphic sites increases power for both window sizes:** Each panel illustrates G12's power to detect hard sweeps for two different window sizes: 51 (orange) and 101 (gold) SNPs . Power was calculated for 200 simulations run for each population (columns) under three different selection coefficients (rows; *s* = 0.01, 0.02, 0.1), continuous (triangle, dashed lines) and non-continuous (circle, solid lines) selection, and three different start times. Grey boxes indicate that the sweep did not occur in that population.

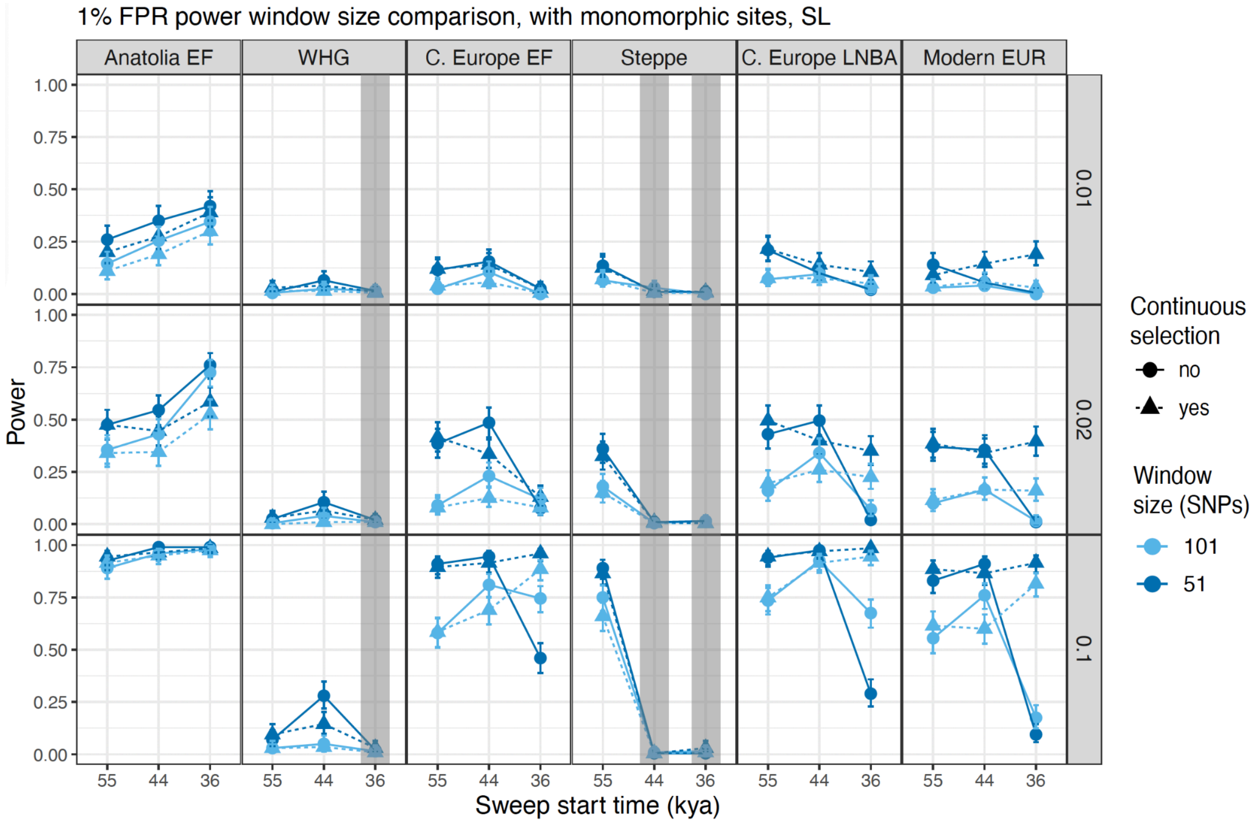


**SI Figure 3 Smaller window sizes tend to increase SL's power to detect sweeps:** Each panel illustrates SL's power to detect hard sweeps for two different window sizes: 51 (dark blue) and 101 (light blue) SNPs . Power was calculated for 200 simulations run for each population (columns) under three different selection coefficients (rows; *s* = 0.01, 0.02, 0.1), continuous (triangle, dashed lines) and non-continuous (circle, solid lines) selection, and three different start times. Grey boxes indicate that the sweep did not occur in that population.

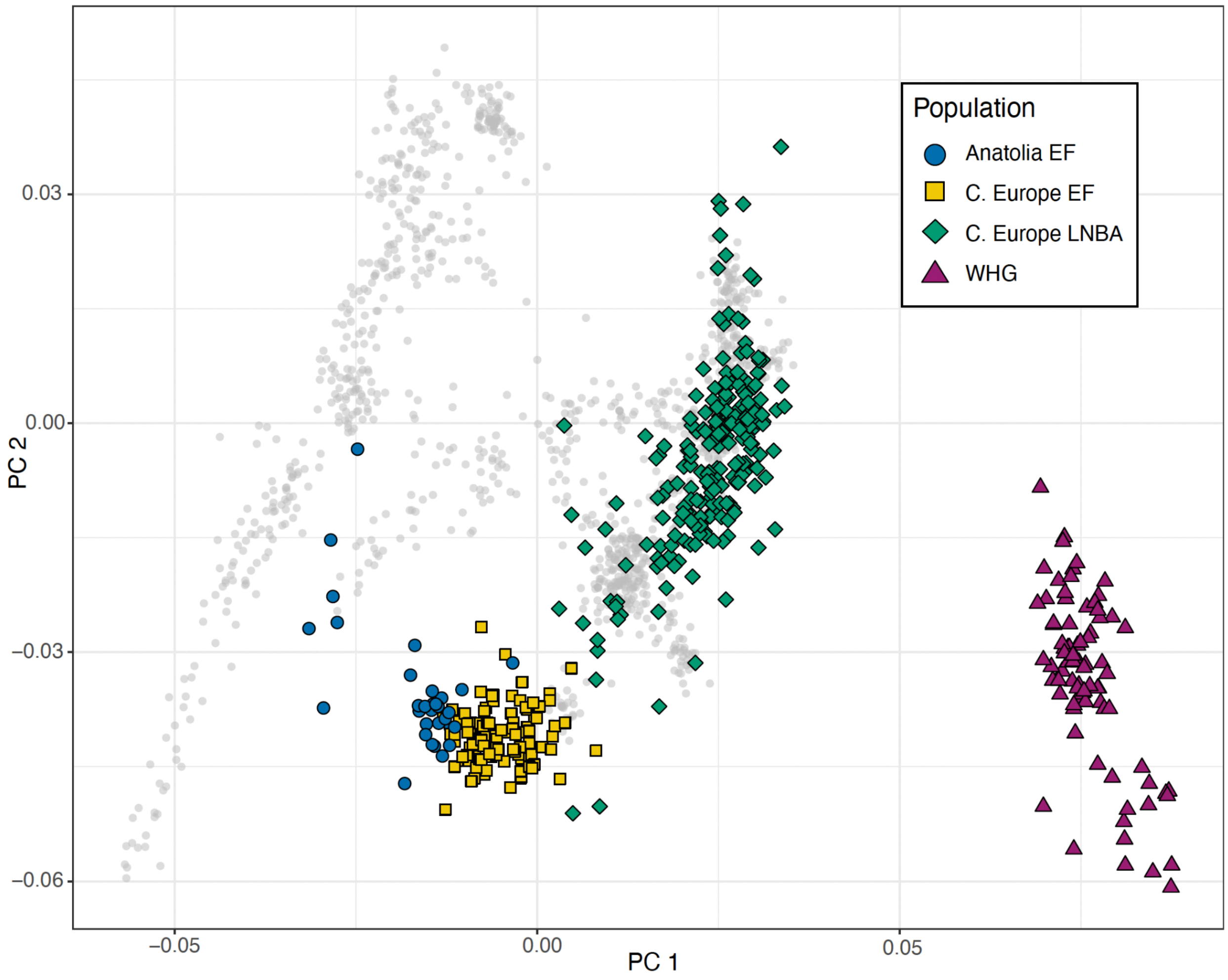


**SI Figure 4 PCA of ancient empirical data:** Grouping was based on the age and geographic location of the sample. Colors and shapes indicated the population group. Grey dots are modern samples from the Human Origins data set.

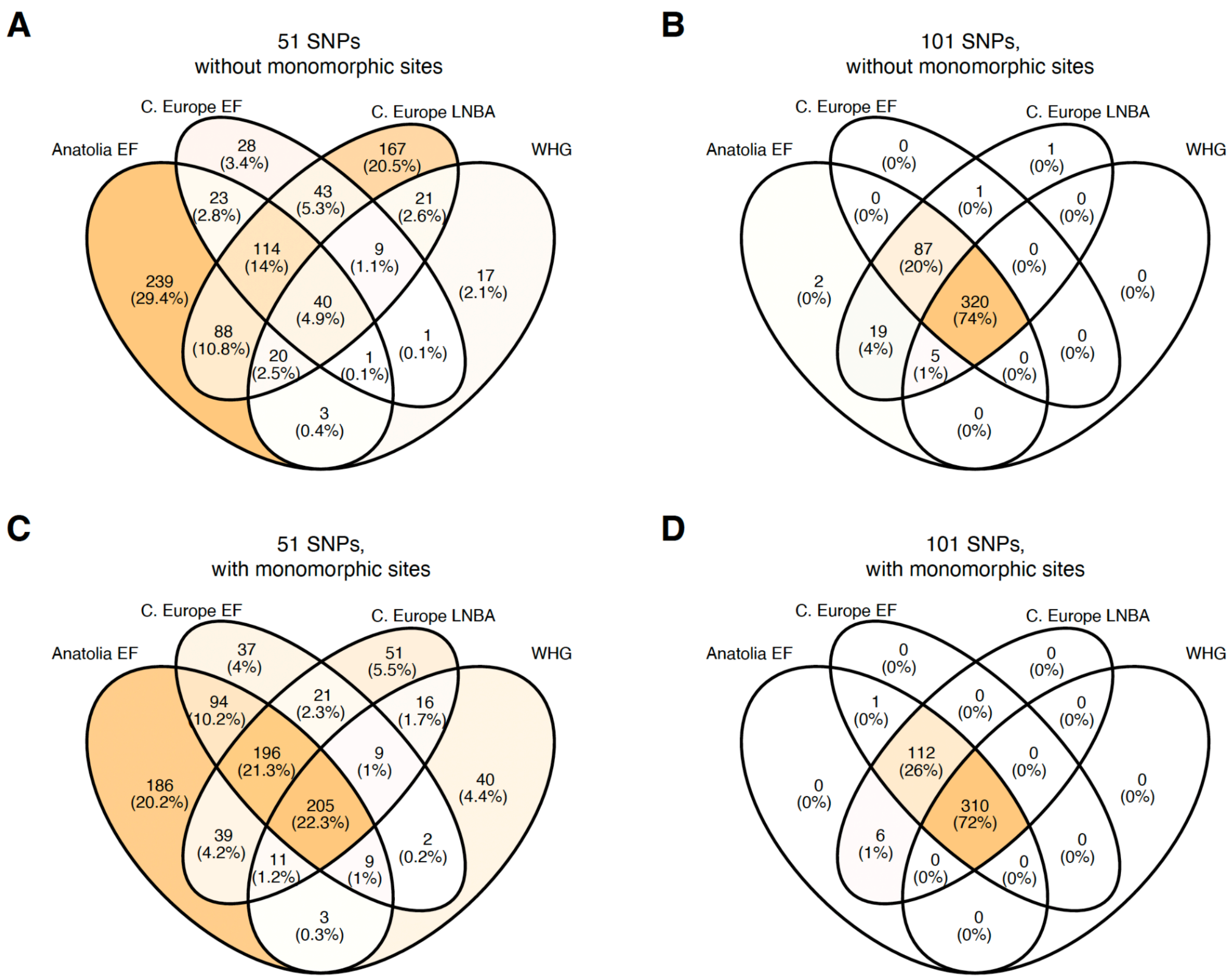


**SI Figure 5 Venn diagram of G12 sweeps shared between empirical populations for *r* = 1 x 10$^{-8}$:** Panels A-D show the total number of sweeps identified and shared between four ancient populations: WHG, Anatolia EF, central European EF and central European Bronze Age. These sweeps were identified with a neutral cutoff computed from neutral simulations run with a recombination rate of 1 x 10$^{-8}$. Panels are broken down by method parameters of window size (51 or 101 SNPs) and whether or not monomorphic sites were filtered out.

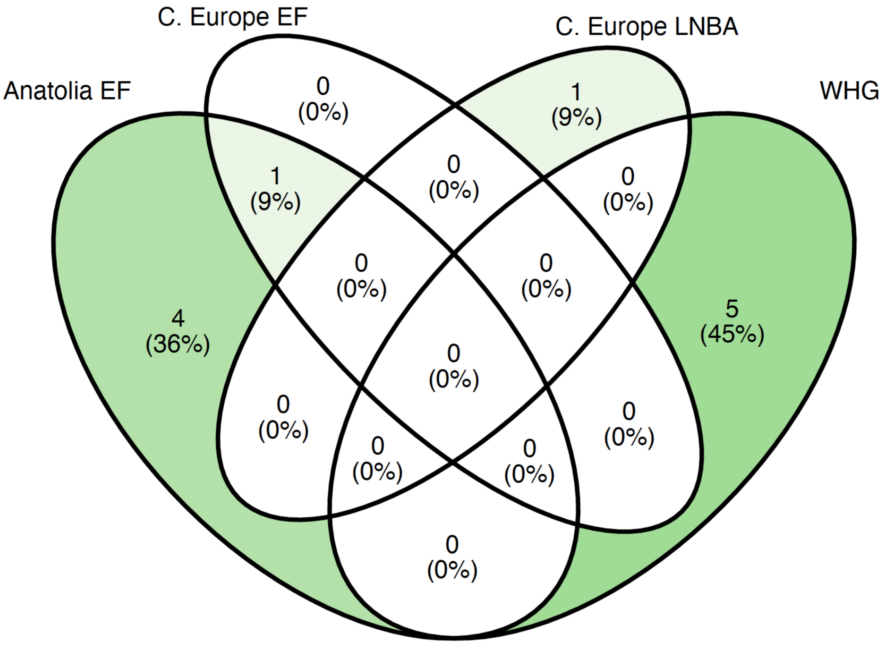


**SI Figure 6 Venn diagram of SF2 sweeps shared between empirical populations for *r* = 5 x $10^{-9}$:** The total number of sweeps identified and shared between four ancient populations: WHG, Anatolia EF, central European EF and central European Bronze Age. These sweeps were identified with a neutral cutoff computed from neutral simulations run with a recombination rate of 5 x $10^{-9}$.

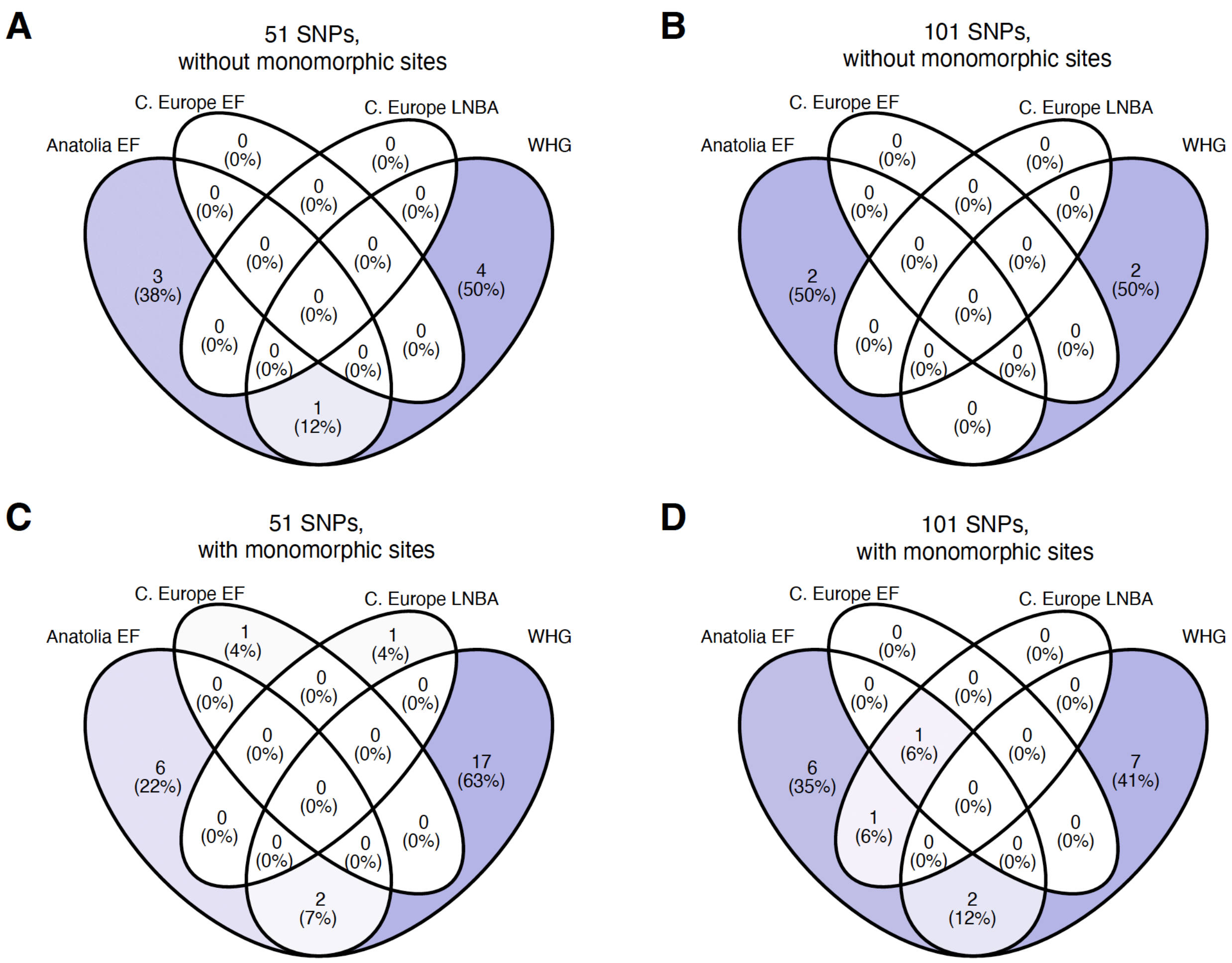


**SI Figure 7 Venn diagram of SL sweeps shared between empirical populations for $r = 5 \times 10^{-9}$:** Panels A-D show the total number of sweeps identified and shared between four ancient populations: WHG, Anatolia EF, central European EF and central European Bronze Age. These sweeps were identified with a neutral cutoff computed from neutral simulations run with a recombination rate of $5 \times 10^{-9}$. Panels are broken down by method parameters of window size (51 or 101 SNPs) and whether or not monomorphic sites were filtered out.

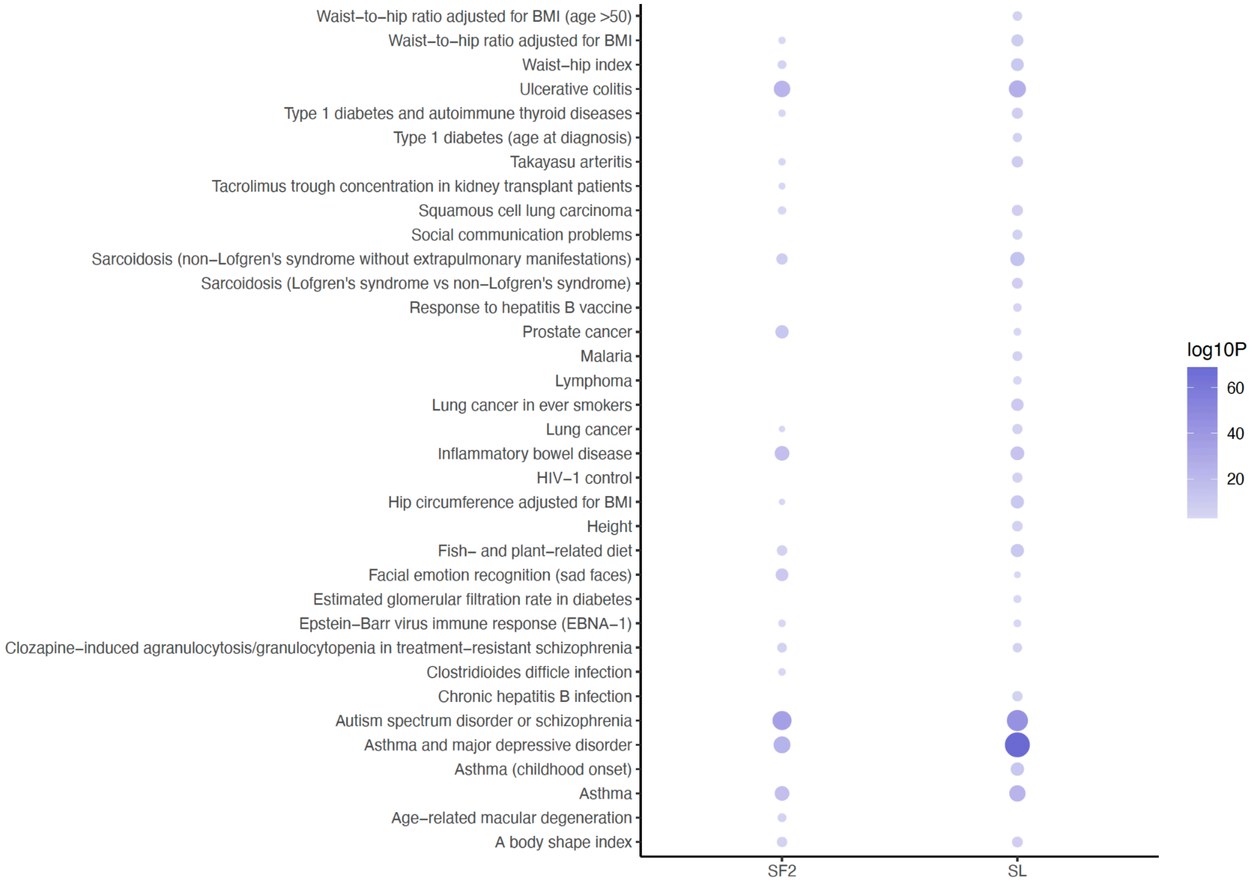


**SI Figure 8 Significantly enriched GWAS categories by method:** FUMA was run on genes that fell within sweep regions for SF2 and SL. We log-transformed the adjusted p-values and considered any category that had above a log value of 3 to be significantly enriched.

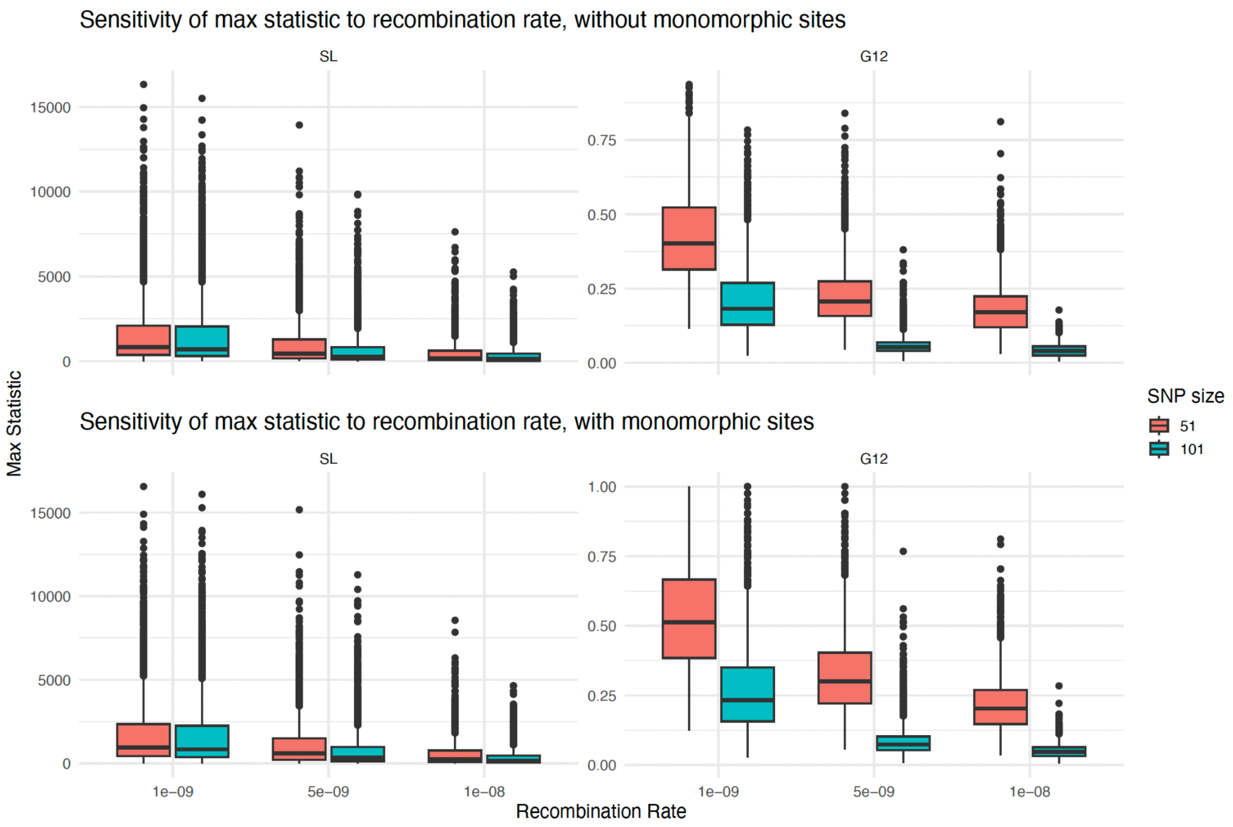


**SI Figure 9 Boxplot of the max statistic for different recombination rates:** The max statistic for each simulation run under three different recombination rates ($1x10^{-9}$, $5x10^{-9}$, and $1x10^{-8}$). Top row illustrates the maximum statistics calculated by SL and G12 for filtered data and the bottom row for the unfiltered data. Left panels represent the values from SL and right panels for G12. The boxplots are colored by window size parameter (red for 51 and blue for 101 SNPs).

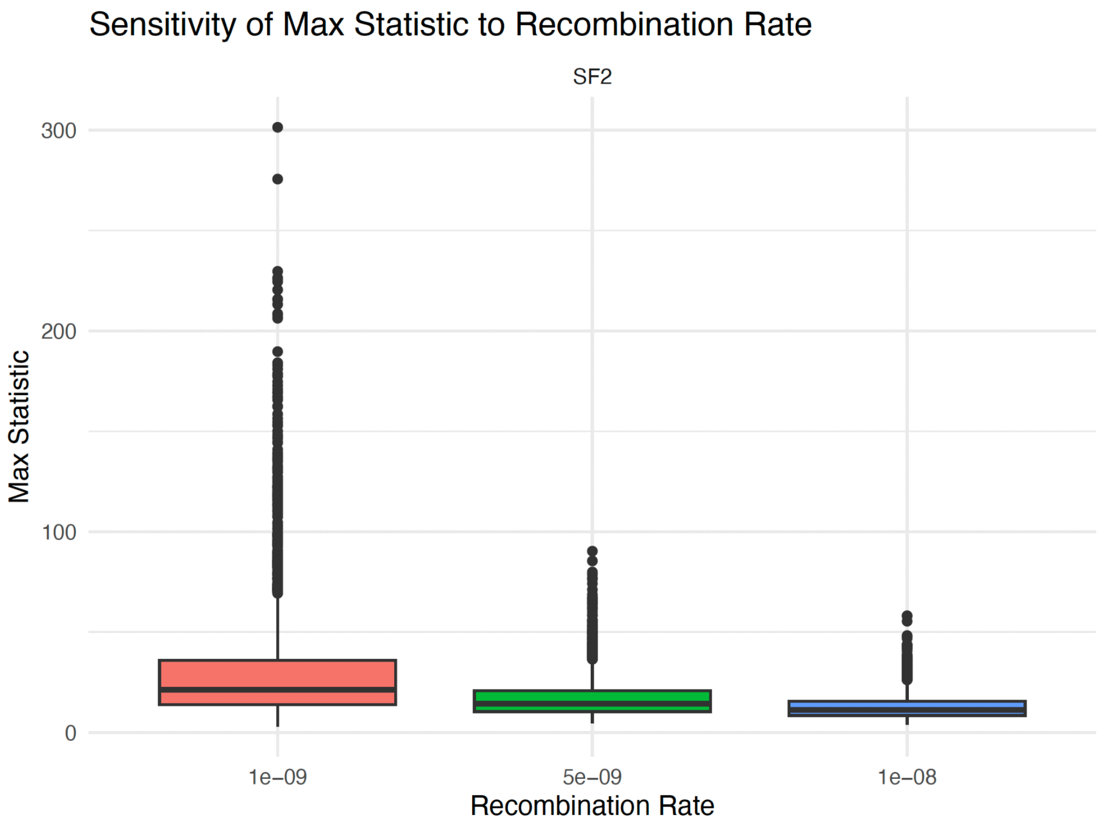


**SI Figure 10 Boxplot of the max statistic for different recombination rates:** The max statistic calculated by SF2 for each simulation run under three different recombination rates ($1x10^{-9}$, $5x10^{-9}$, and $1x10^{-8}$).
